# Heme induces innate immune memory

**DOI:** 10.1101/2019.12.12.874578

**Authors:** Elisa Jentho, Boris Novakovic, Cristian Ruiz-Moreno, Ioannis Kourtzelis, Rui Martins, Triantafyllos Chavakis, Miguel Perreira Soares, Lydia Kalafati, Joel Guerra, Franziska Roestel, Peter Bohm, Maren Godmann, Tatyana Grinenko, Anne Eugster, Martina Beretta, Leo A.B. Joosten, Mihai G. Netea, Hendrik G. Stunnenberg, Michael Bauer, Sebastian Weis

## Abstract

Trained immunity defines long-lasting adaptive responses of innate immunity, mediated by transcriptional and epigenetic modifications of myeloid cells. Here, we report that labile heme, an alarmin released extracellularily upon tissue damage and hemolysis, induces trained immunity in human and murine primary cells and in mice. Heme-trainning in monocytes is associated with epigenetic and transcriptional profiles that overlap to some extent with those seen in β-glucan-training, the prototype agonist of trained immunity. In sharp contrast to β-glucan training however, heme-training relied on activation of the Syk/JNK-pathway. Heme training in mice was associated with long-lasting expansion of myeloid-biased progenitor cells as well as with altered chromatin accessibility in hematopoietic stem and progenitor cells. Finally, heme-induced training was protective against bacterial sepsis in mice.

In conclusion, we reveal that heme, a *bona fide* alarmin, induces innate immune memory regulating host response to infection.

## INTRODUCTION

Heme is an iron-containing protoporphyrin that acts as a prosthetic group in vital hemoproteins^1^. Hemolysis or other forms of tissue damage can cause accumulation of cell-free labile heme in plasma ^2^. As it becomes loosely bound to plasma acceptor proteins or macromolecules, these reduce but do not prevent labile heme redox activity that underlies its pathological effects ^2,3^. As such, labile heme acts as a highly pro-oxidant agonists that contributes critically to the pathogenesis of severe infections, as demonstrated for malaria ^4–7^ and for bacterial sepsis ^8–10^. Labile heme can be considered a prototypical alarmin ^9^. It is an intracellular molecule that is released upon cellular damage and sensed by innate immune cells via different pattern recognition receptors (PRR), including Toll Like Receptor (TLR)-4 and the NOD-like receptor family ^11,12^. It also induces inflammasome activation in murine macrophages (Mφ) when applied shortly after lipopolysaccharide (LPS) ^13^. Of note, chronic low-dose cellular exposure to labile heme can exert protective effects via the induction of the heme catabolizing enzyme heme oxygenase-1 (HO-1), underlying exemplarily how sickle cell trait protects against malaria ^6^. Also depending on dosage and timing of administration, heme can exert protective effects against a variety of other immune-mediated inflammatory diseases ^14^. It is now well established that certain pathogen associated molecular patterns (PAMPs), such as LPS, fungal β-glucan or certain vaccines can induce long-lasting epigenetic and transcriptional modifications in myeloid cells ^15–18^. This is perceived as part of an evolutionarily conserved host defense strategy against infections caused by different pathogens ^18–20^. The deposition of epigenetic marks on histone tails, poising target genes for transcriptional activation or repression in response to PRR signaling by PAMPs ^18,19^. This contributes to the long-known phenomenon of LPS tolerance ^15,21^. However, depending on the stimulating component, it can also result in an enhanced inflammatory response upon restimulation which is then referred to as trained immunity or innate immune memory. Trained immunity was originally demonstrated to occur in monocytes and Mφ and later on in other cell types, such as Natural Killer (NK) cells ^22,23^ or epithelial stem cells ^24^. It can be initiated at the level of hematopoietic stem and progenitor cells (HSPC), associated with increased myelopoiesis ^25,26^.

While PRR likely evolved to recognize PAMPs expressed by microbes ^27^, these can also sense a number of endogenous molecules, designated originally as damage-associated molecular patterns (DAMP)^28,29^ or alarmins ^9^. Alarmins can signal via PRR as well as other receptors to promote inflammatory responses. In fact, the mutual inflammatory response to PAMP or DAMP is preserved independently of the presence of a given pathogen ^29^. If so, secondary responses to primary inflammatory cues should be conserved between PAMP and DAMP ^30^.

Here, we report that labile heme^9^ induces potent trained immunity *in vitro* in human monocytes and Mφ, as well as *in vivo* in mice. We identified epigenetic and transcriptional overlaps as well as distinct differences between the monocyte response to heme and to β-glucan, the prototype inducer of trained immunity. Heme training relies on Spleen tyrosine kinase (Syk) and c-Jun n-terminal kinase (JNK) activation, but not on the activation of the Mammalian Target of Rapamycin (mTOR), with the latter being β-glucan training ^31^. Heme training is also associated with the expansion of bone marrow HSPCs towards the myeloid linage and long-term chromatin accessibility patterns. Finally, heme-training is protective in polymicrobial sepsis, simailr to other types of trained immunity.

## RESULTS

### Heme induces innate immune memory

We hypothesized that beyond its acute inflammatory properties ^12,32^, labile heme can induce long-term memory in myeloid cells, *i.e.* innate immune training. Using a well established 2-hit model (*Figure (Fig.) 1a*) ^33^ we found that heme increased cytokine release from human and murine primary Mφ upon secondary challenge by LPS (*Fig. 1b,c; Supplementary (Suppl.).Fig1*), consistent with the notion of heme affording trained immunity.

**Figure 1:**
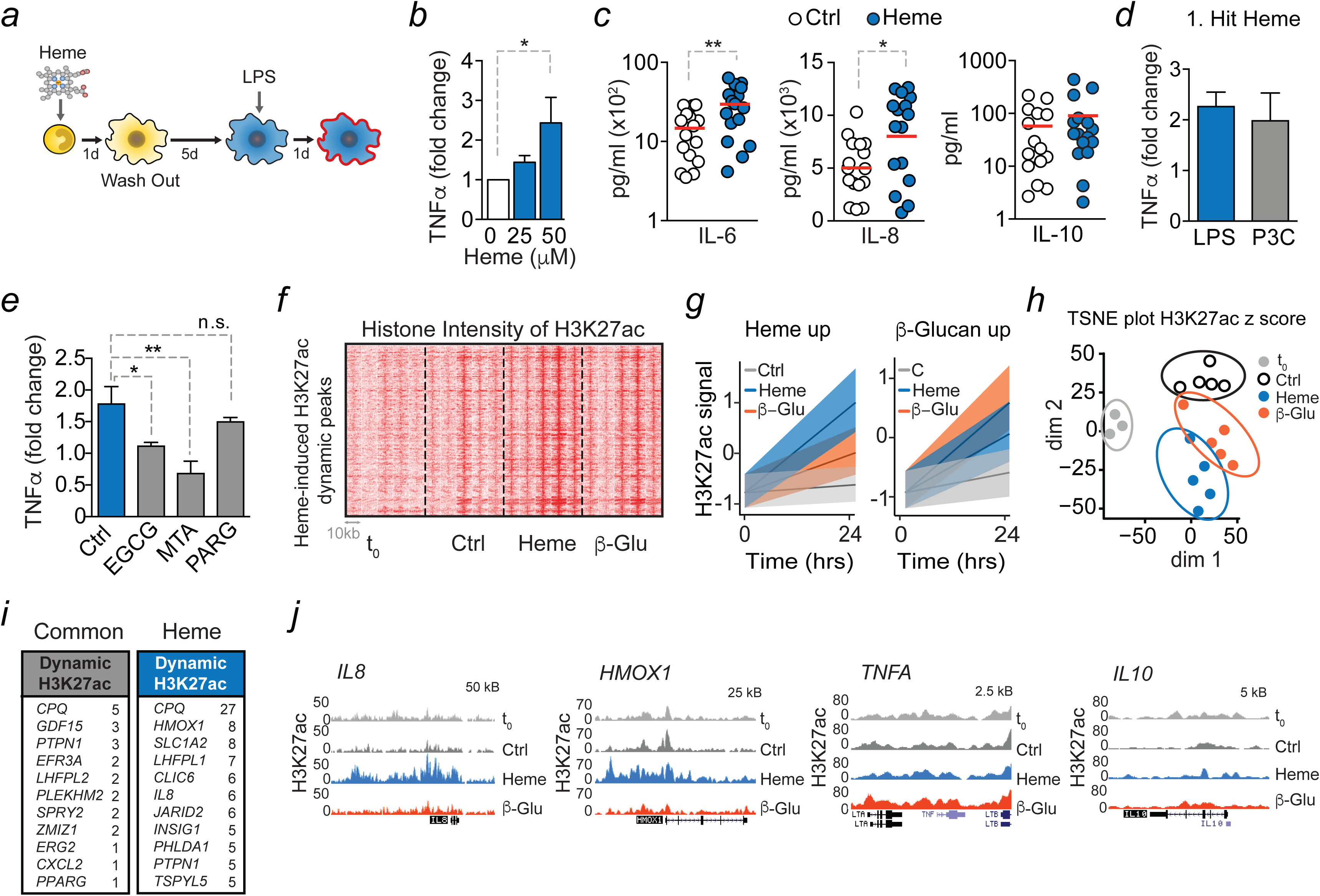
Heme imposes innate immune training. **(a)** Experimental setup. **(b-c)** Cytokine release from vehicle or heme-trained Mφ after LPS application (b) TNF measurements by ELISA (c) IL-6 and IL-10 measurements by LegendPlex™. IL-8 measurement by ELISA. n=16 independent donors. **(d)** TNF release from vehicle or heme trained Mφ, 2^nd^ hit: LPS or Pam3CSK4. **(e)** TNF release from vehicle or heme trained Mφ treated with respective inhibitors. **(f)** Heatmap of H3K27ac intensity at 497 regions with 5 kb ± from center of the peak. **(g)** H3K27ac intensity of β-glucan and heme specific regulated acetylation marks at baseline and 24 h after training. Black line is median z-score and shaded areas are 25% and 75% confidence intervals. **(h)** TSNE-plot of dynamic acetylation peaks in monocytes and vehicle, β-glucan or heme trained Mφ. Dim1: differentiation processes. Dim2: variation due to treatment. **(i)** Genes with highest number of differential H3K27ac peaks within 1 Mb of their transcription start site. **(j)** Representative gene loci corresponding to *IL8*, *HMOX1*, *TNFA* and *IL10*. For all epigenetic analysis n=5 individual donors. n=3 independent experiments à 2-3 donors per experiment unless indicated otherwise. All data mean±SEM unless otherwise stated. Student’s t-test or One-way ANOVA with Fisher’s LSD. *p≤0.05; **p≤0.01; ***p≤0.001. <u>Abbreviations:</u> β-Glu.. β-glucan; Ctrl.. control; d.. days; EGCG.. epigallocatechin-3-gallate; hrs.. hours; H3K27ac.. histone 3 lysine 27 acetylation; HMOX1.. Heme oxygenase 1; IL.. Interleukin; LPS.. lipopolysaccharid; MTA.. 5′-deoxy-5′-methylthioadenosine; n.s…non-significant; PARG.. pargyline; P3C.. Pam3CSK4; t_0_.. baseline before stimulation; dim.. dimension; TNF.. Tumor Necrosis Factor; TSNE.. t-Distributed Stochastic Neighbor Embedding.

The increased cytokine release was non-specific to TLR-4 signaling as activation of TLR1/2 signaling by the synthetic receptor agonist Pam3CSK4 used as the secondary challenge, also resulting in increased TNF-release (*Fig. 1d*).

Training effects were specific to heme as *i*) different concentrations of LPS as a first stimulus resulted in immune tolerance and *ii)* neither the application of ATP as a first hit nor heme as a second hit altered TNF release (*Suppl.Fig. 1*). Of note, while most of the effects of heme have previously been associated with its pro-oxidative properties imposed by the central iron atom, this is not the case for heme training. We find that it can also be induced by protoporphyrin (PPIX), lacking iron (*Suppl.Fig.1*). Accordingly, heme-induced trained immunity was not prevented by the glutathione precursor N-Acetyl Cysteine (NAC) (*Suppl.Fig.1*), which counteracts heme/iron-driven oxidative stress ^10^.

Priming of innate immune cells by β-glucan was linked to transcriptional and epigenetic changes, *such as* acetylation of histone 3 lysine 27 (H3K27ac) ^16–19^. Accordingly, heme-induced training was significantly decreaed by acetylation and methylation inhibitors (*Fig. 1e*). We then performed ChIP-sequencing in untreated monocytes and 24 hours after exposure to 50 μM heme ^34^ in order to assess the epigenetic profile of heme-induced training. In this set of experiments, we included β-glucan, the prototype trained immunity agonist, as a positive control allowing the identification of common trained-immunity related chromatin remodeling events regardless of the primary stimulus. After heme exposure, approximately 5% of all H3K27ac marked regions in the genome showed increased signal as compared to untreated controls (*Fig. 1f; Suppl.Fig. 2*). Induced H3K27ac regions could be separated into heme-specific and heme/β-glucan common (*Fig. 1g; Suppl.Fig. 2; Suppl.Tables 1;2*). The t-SNE plot (dimension 2) clearly separated heme and β-glucan-trained and naïve Mφ by H3K27ac pattern (*Fig. 1h*). Overall, 24 h after heme application, 497 peaks in the proximity of 308 genes showed increased H3K27ac signal. Dynamic H3K27ac regions were assigned to the nearest gene, with several genes having two or more regions in their proximity (*Fig. 1i*). Examples of H3K27ac tracks for genes of interest are shown in *Figure 1j*. H3K27ac dynamics are observed at both promoters and distal enhancer regions, with 98.6% of peaks occurring within 0.5Mb of a gene TSS (*Suppl.Fig. 2*). Gene ontology (GO) analysis showed that heme exposure elevated H3K27ac signal near genes involved in wounding and stress, which is in accordance with the notion that heme acts as a DAMP. In contrast, H3K27ac levels at genes related to adaptive immune regulation were down-regulated by heme (*Suppl.Fig. 2*).

**Figure 2:**
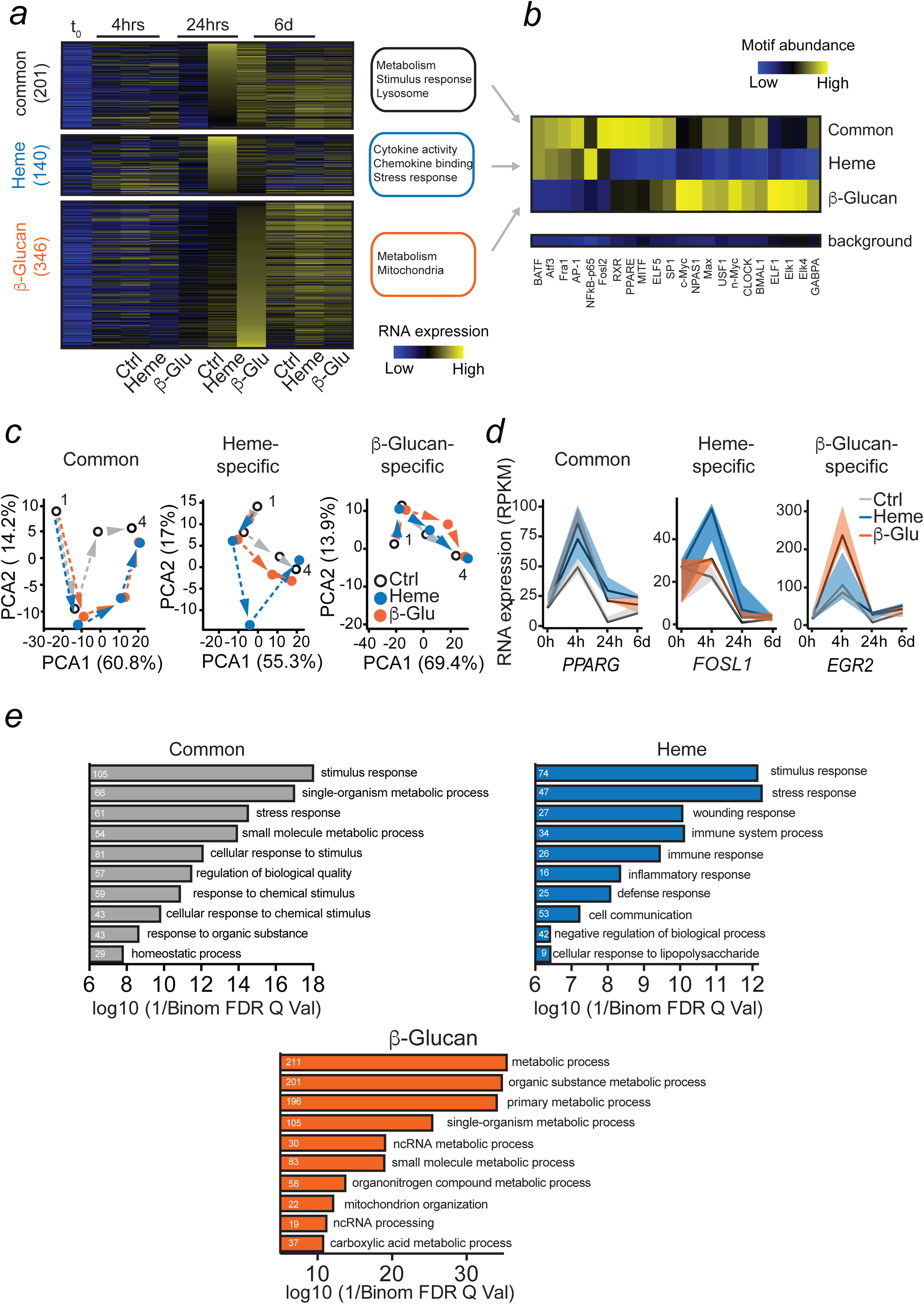
Heme training evokes a specific transcriptional signature. Transcriptional analysis by RNAseq of control and heme or β-glucan trained myeloid cells at various time points after the first hit. **(a)** Heatmap of regulated genes at baseline (t_0_) and at indicated time points after heme or β-glucan exposure. Up-regulated genes were grouped in commonly-regulated (similar regulation in β-glucan and heme treated cells) and heme or β-glucan-specifically regulated. Gene ontology terms are listed next to each group of genes. **(b)** Heatmap of respective transcription factor motifs enrichment (% abundance) at promoters of commonly, heme- or β-glucan-specifically regulated genes 24 hrs after heme/β-glucan. **(c)** PCA plot of the dynamic gene expression of commonly, heme- or β-glucan-specifically regulated genes at different time points. PC1 is primarily explained by differentiation, while PC2 represents treatment. Dots represent different time points: 1=t_0_, 4=6 days. **(d)** Time-resolved expression patterns of selected transcription factors that are both induced by Heme and/or β-glucan and have an enriched motif at the promoters of downstream genes shown in ***b**.* **(e)** Most abundant pathways based on gene expression analysis 24 h after treatment. White numbers indicate number of affected genes. All analysis were performed from n=5 individual donors. <u>Abbreviations:</u> β-Glu.. β-glucan; Ctrl.. control; hrs.. hours; d.. days.

### Heme induces distinct transcripts and common markers of innate immune memory

We then asked whether the transcriptome of heme-induced trained *vs.* non-heme trained cells differs over time (*Fig. 2*). RNA sequencing (RNA-seq) of human monocytes and Mφ treated with vehicle, heme or β-glucan was performed at different time points. The largest number of differentially expressed genes occurred at 24 hours after stimulation, with 201 genes commonly induced by heme and β-glucan, 140 specifically induced by heme and 346 specifically induced by β-glucan (*Fig. 2a; Suppl.Table 3;4;5*). The common signature was related to *lysosome maturation* and *metabolism*, key processes required for increased cytokine production observed in trained Mφ. Genes only induced by heme are mainly involved in inflammatory pathways (*Fig. 2a*). Motif analysis of the gene promoter regions indicated that there is a β-glucan-specific signature (recognised by: *USF* and ELK1), heme-specific signature (*e.g.* NF-κB, BATF) and a common signature (AP-1, PPAR and MITF) (*Fig. 2b*). Principial component analysis (PCA) plots based on the expression of common genes showed that heme and β-glucan exposed cells move faster along the differentiation trajectory (*Fig. 2c*), as previously reported for β-glucan ^34^. We were also interested whether heme-induced trained immunity was associated with a polarization into classical M1, M2 ^35^ or hemorrhage-associated Mφ (M-hem) ^36^. Yet, using transcriptomic data from different time points, heme-trained Mφ did not display any of this typical polarization traits (*Suppl.Fig. 2*). Consistent with the AP-1 motif enrichment, transcription factors components of AP-1, namely *JUN* and *FOSL* were induced by heme, while *ELK1* mRNA was induced exclusively by β-glucan (*Suppl.Fig. 3*). We also observed the expected upregulation of Dectin-1 target *EGR2* by β-glucan exposure and upregulation of *MITF* and *PPARG* by both β-glucan and heme (*Fig. 2d; Suppl.Fig. 3*). Heme application specifically induced transcription of heme/iron related genes *HMOX1, FTH1, HRG1* but not of the iron exporter ferroportin (*FPN1*) (*Suppl.Fig. 3*). Pathway analysis using GREAT ^37^ demonstrated that those pathways involved in cytokine-signaling and stress-response were up-regulated during heme training, in contrast to genes encoding metabolism and mitochondrial function in β-glucan training. Common pathways included metabolic and lysosome regulation (*Fig. 2e*), indicating that these pathways may be fundamental for innate immune memory. Gene set enrichment analysis (GSEA) using the Molecular Signatures Database (MSigDB) (*Fig. 2e*) showed that common genes between heme and β-glucan induced pathways are involved in stress response, while the heme-specific response is related to inflammation and immune response and β-glucan induced pathways relate rather to metabolic processes. This is in line with the motif analysis, indicating that heme induces one set of inflammatory genes, and another set of trained immunity-inducing genes, being shared with β-glucan. Heme-trained bone marrow-derived macrophages (BMDM) from C57Bl/6 mice increased HO-1 protein content after the 2^nd^ hit supporting our finding in human cells that heme induces epigenetic modifications of the *HMOX1* promoter (*Suppl. Fig. 1*).

**Figure 3:**
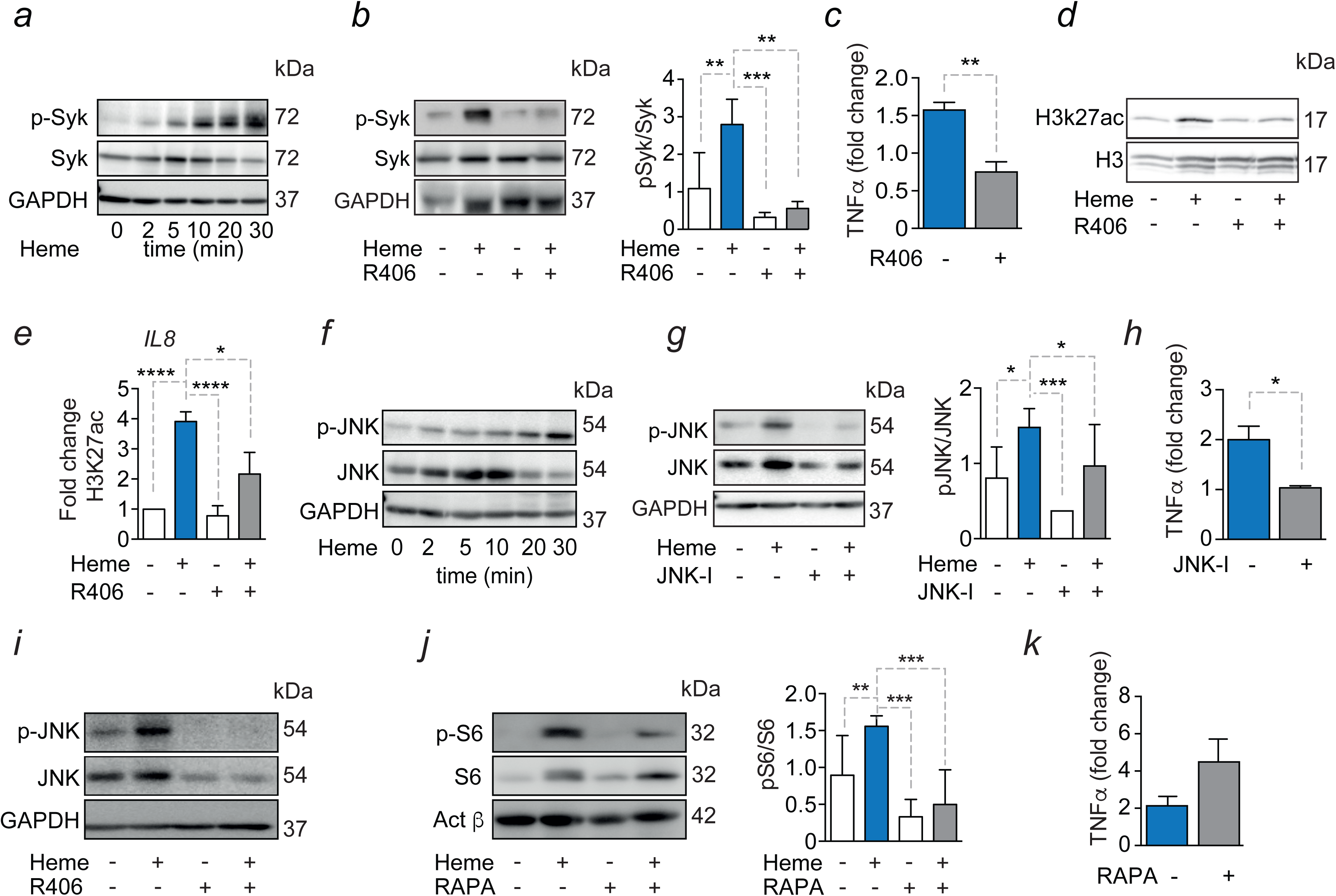
Heme training relies on Syk/JNK activation. (**a**) Western blot of Syk-phosphorylation in human monocytes after heme treatment. **(b)** Syk-phosphorylation in human monocytes 20 min after heme stimulation ± the Syk-inhibitor R406. (**c**) TNF production after re-stimulation with LPS of vehicle or heme trained Mφ ± R406. (**d**) Western blot of total H3K27ac 24 h after heme treatment of human monocytes ± R406. (**e**) ChIP-qPCR analysis of H3K27ac at the *IL8* promoter (n=3 independent experiments). (**f**) Western blot of JNK-phosphorylation in human monocytes after heme treatment. **(g)** Western blot and densitometry of JNK activation in human monocytes 20 min after heme stimulation ± the JNK-inhibitor SP600125. (**h**) TNF production after re-stimulation with LPS of vehicle or heme-trained Mφ ± SP600125. (**i**) Western blot of JNK phosphorylation in human monocytes 20 min after heme stimulation ± R406. **(j)** Western blot of p-S6 in human monocytes 24 h after heme stimulation. **(k)** TNF production after re-stimulation with LPS of vehicle or heme-trained Mφ ± rapamycin. TNF values normalized to LPS controls or cells reciving inhibitor and LPS. Representative Western blot of at least 3 independent donors. Densitometries were analyzed by One-way ANOVA with Fisher’s LSD; mean ± SEM. All ELISA results derived from 3 independent experiment à 3 donors each and were analyzed by Students t-test; mean± SEM; *p≤0.05; ** p≤0.01; ***p≤0.001; ****p≤0.0001. <u>Abbreviations:</u> GAPDH.. Glyceraldehyde 3-phosphate dehydrogenase; H3K27ac.. histone 3 lysine 27 acetylationJNK.. c-Jun-N-terminal kinases; JNK-I.. JNK-Inhibitor SP600125; LPS.. Lipopolysaccharide;; Syk.. Spleen tyrosine kinase; TNF.. Tumor necrosis factor.

### Heme training is mediated via a Syk/JNK dependent pathway

Having established that heme induces innate immune memory via histone modifications, we were interested in the underlying mechanism. Consitent with previous reports^32^, heme appears to signal via the spleen tyrosine kinase (Syk), as assessed by its phosphorylation in human monocytes (*Fig. 3a; Suppl.Fig.3*). Moreover, Syk inhibition by R406 ^38^ abolished heme-induced training (*Fig.3b;c*). Global increase in H3K27ac in response to heme treatment was also prevented by Syk-inhibition, as assessed by Western bBlot in human monocytes (*Fig.3d*). Specifically, ChIP-qPCR analysis of the *IL8* promoter showed that heme-induced H3K27ac deposition was prevented by R406, pointing towards an association of heme-induced Syk activation and Histone 3 acetylation (*Fig.3e*). Moreover, ChIP-qPCR analyses confirmed the ChIP-seq findings as a locus-specific increase in H3K27ac after heme training could be detected in promotor regions of selected genes, (*e.g. IL8)* which was reduced upon inhibition of Syk signaling (*Fig. 3e*) in line with our previous findings.

In keeping with the notion that JNK can act down stream of Syk activation ^13^, we assessed whether heme-induced training relies on JNK activation. Heme exposure rapidly increased JNK-phosphorylation in human monocytes (*Fig. 3f; Suppl.Fig. 3*) while application of the JNK-Inhibitor SP600125 ^39^ abolished JNK phosphorylation and heme-induced training (*Fig. 3g;h*). In further support of this notion, the Syk-inhibitor R406 decreased JNK phosphorylation, suggesting that heme-induced training is mediated via Syk-dependent JNK-activation (*Fig. 3i*).

We also assessed whether mTOR signaling, which mediates β-glucan training, was required for heme training as well ^31^. Despite increased phosphorylation of the mTOR target S6 in heme-trained Mφ (*Fig. 3j*), inhibition of mTOR activity by rapamycin did not abolish heme training, as the increased TNF production of heme trained cells to a secondary challenge was not affected by the mTOR inhibitor (*Fig. 3k*). Consistently, application of the Syk-inhibitor R406 during heme training did not alter the S6-phosphorylation indicating that activation of mTOR by heme acts independently of Syk signaling (*Suppl.Fig. 3*). Heme-training was independent of TLR4 signaling, previously shown to be induced by heme ^11^ as BMDM from C57Bl/6 mice with a constitutive deletion of *Tlr4* (*Tlr4^−/−^*) were equally trained by heme, using Pam-3-Cys as the second stimulus (*Suppl.Fig. 1*).

### Heme training promotes myelopoiesis

Modulation of myelopoiesis progenitors is an integral component of innate immune training ^25,26^. Seven days after heme administration, we found no difference in cell number or percentages of LSK cells (Lin^−^cKit^+^Sca1^+^), long-term (LT)-HSCs (CD48^−^CD150^+^LSK), or multipotent progenitors (MPP, CD48^+^CD150^−^LSK), as compared to control-treated mice. However, we observed increased frequency of myeloid-biased LT-HSCs (CD41^+^ LT-HSC) ^40^ amongst the LT-HSC population as well as of myeloid-biased MPP3 cells (Flt3^−^CD48^+^CD150^−^ LSK) with a corresponding decrease of lymphoid-biased MPP4 cells (Flt3^+^CD48^+^CD150^−^ LSK) (*Fig. 4a,b*). Myeloid-biased LT-HSCs (CD41^+^ LT-HSC) and MPP3 were still increased 28 days after heme treatment (*Suppl.Fig. 4*). Furthermore, a significant increase of mature neutrophils and monocyte is observed at this time point. Given that long-term adaptations would require changes in the epigenome landscape, we assessed genome-wide chromatin accessibility using single nuclei ATAC sequencing on purified LSK cells at seven days (Control = 4,842 cells and heme = 5,562 cells) and 28 days (Control = 6,762 cells and heme = 6,052 cells) after heme training (*Fig. 4c-f, Suppl.Fig. 5,6*). We identified eight cell subpopulations (*Fig. 4c*) based on two approaches: first, matching known bulk ATAC profiles of the mouse immune system ^41^ and second, creating gene activity scores which allow the exploration of specific gene markers ^42,43^ ^44^ and plot their distribution over the cell clusters. Within the HSC/MPP3-4 cluster, we found a transition of progenitors from LT-HSC towards MPP3 and MMP4 (*Fig.4d*). After seven days of heme exposure, there was a modest increase in the proportion of HSC-MPP3-4 cells compared with control mice (*Fig.4e,f*). Strikingly, upon 28 days of heme treatment, several cell subpopulations were diminished, particularly MEP, and cells mostly concentrated on the HSC-MMP3-4 cluster (*Fig. 4g,h; Suppl. Fig. 5,6*). Of note, we identified a group of cells (refered to as enriched MPP (eMPP)) within the earliest progenitors with a low LT-HSC signature score but enriched for MPP (*Fig. 4g,h*).

**Figure 4:**
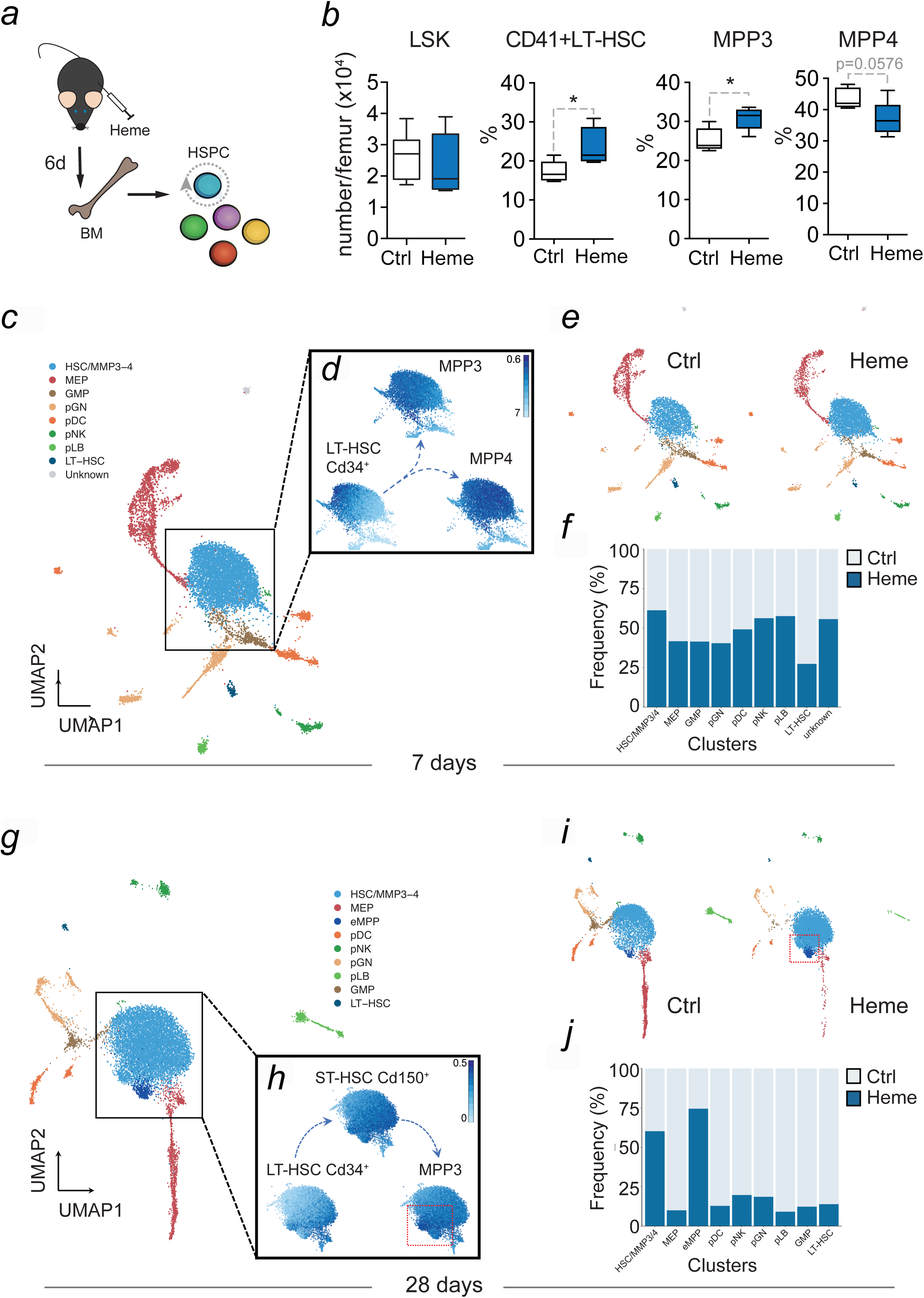
Heme training expands myeloid-biased hematopoetic stem cells in mice. **(a)** Experimental setup. **(b)** Absolute numbers per femur of LSK, MPP and LT-HSC and relative amount of CD41^+^ LT-HSC; MPP3; MPP4 of vehicle or heme-trained mice at day 7. n=6 animals per group derived from 2 independent experiments. Student’s t-test; mean ± SD. *p≤0.05; ***p≤0.001. (**c-f**) Single-nuclei (sn) ATAC analysis from sorted LSK cells after 7 days (or) 28 days of heme treatment. Colored by data-driven cluster number. (**c, g**) Uniform Manifold Approximation and Projection (UMAP) for snATAC-seq data at 7 days (n = 10,404 cells) and 28 days (n= 12,814). Annotation of the clusters was perfomed according to overlaid of epigenetic signatures and known genes (see Materials and Methods) ^41^. (**d, h**) Enrichment of epigenomic signatures (AUC scores) for bulk ATAC-seq profiles of HSC and MPP progenitors from the *ImmGen Database*. (**e, f**) Control and heme UMAP plots side by side and (**i, j**) percentage of contibution per condition to each cluster. <u>Abbreviations:</u> BM.. bone marrow; Ctrl.. Control; HSC.. hematopoietic stem cell; HSC/MPP3-4.. Hematopoetic Stem Cell / Multipotent progenitor; LSK..Lin^−^Sca1^+^cKit^+^; LT.. long term; MPP.. multipotent progenitor cells; MEP.. megakaryocyte erythroid progenitor; GMP.. granulocyte macrophage progenitor; pGN.. precursor of granulocytes; pDC.. precursor of dendritic cells; pNK.. precursor of NK cells; pLB.. Precursor of B lymphocytes; LT-HSC.. Long term Hematopoetic Stem Cell; eMPP.. enriched for MPP cells.

**Figure 5:**
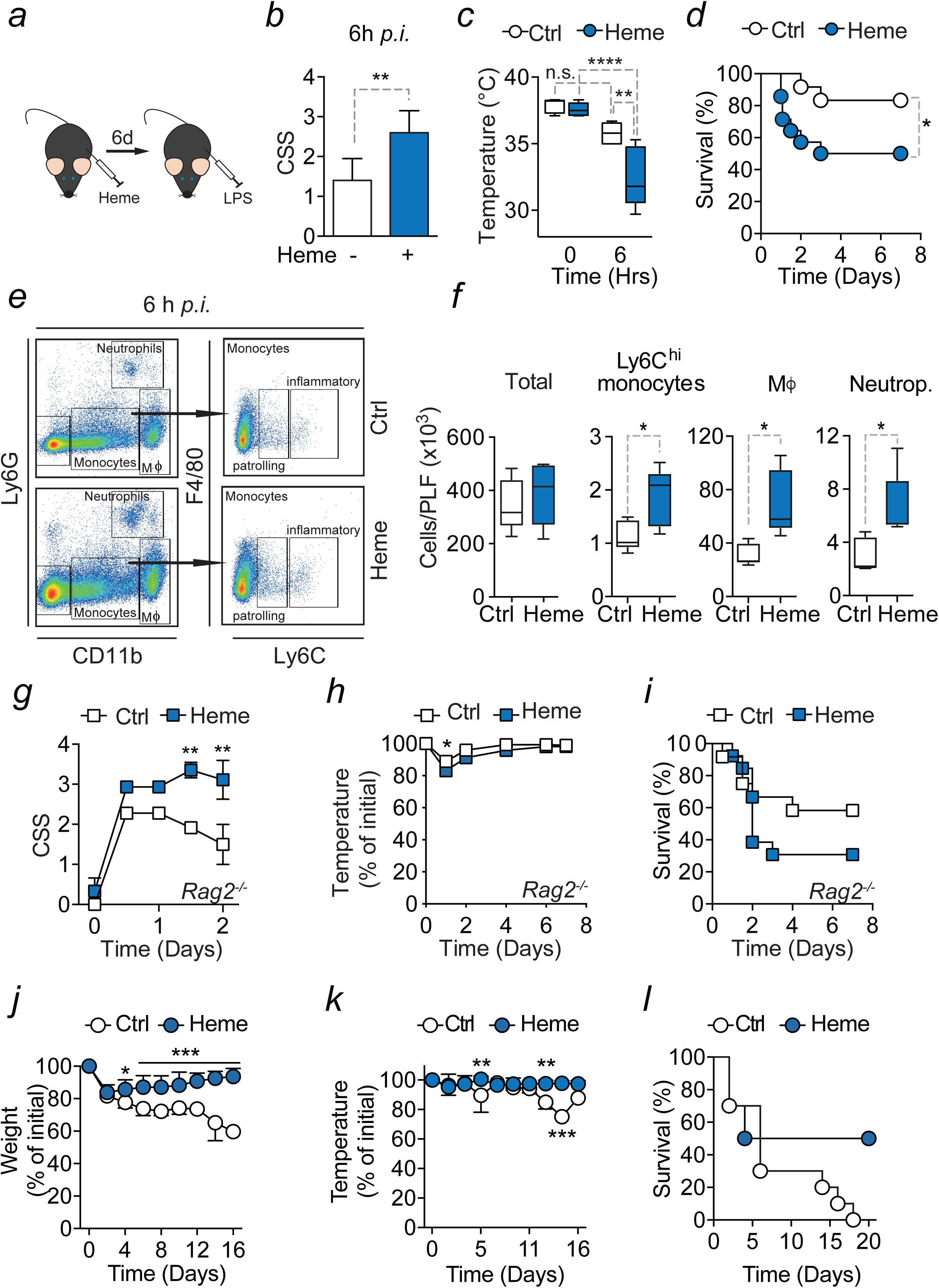
Heme training *in vivo*. (**a**) Experimental setup for heme-induced trained immunity and LPS shock *in vivo.* **(b)** CSS and **(c)** temperature of vehicle or heme-trained mice before and 6 h after *i.p.* LPS treatment. n=5 mice per group; Students t-test or Two-way ANOVA with Fisher’s LSD; mean±SD. **(d)** 7 day survival of vehicle (n=12) or heme-trained (n=14) mice after LPS injection. Pool from 3 independent experiments. **(e)** Scheme of Mφ and monocyte gating strategy. All cells were gated as viable single cells. Neutrophils were gated as CD11b^+^, Ly6G^+^, monocytes were gated as CD11b^+^, Ly6G^_^, and further distinguished as Ly6C^high^ (inflammatory monocytes) and Ly6C^low^ (patrolling monocytes). Peritoneal Mφ were gated as cd11b^high^, Ly6G^_^, F4/80^+^. **(f)** Cell count and flow cytometry analysis of peritoneal lavage fluid. **(g-i)** *Rag2^−/−^* mice subjected to *i.p.* LPS. (g) CSS (h) relative temperature change and (i) survival of vehicle (n=12) and or heme-trained (n=13) *Rag2^−/−^* mice. Pool of 3 independent experiments. **(j,k,l)** Vehicle or heme trained mice subjected to PCI induced sepsis. (j) Weight, (k) temperature and (l) survival of vehicle (n=10) and or heme-trained (n=10) mice. Pool of 2 independent experiments. Two-way ANOVA with Bonferroni; mean±SEM. *p≤0.05; **p≤0.01; ***p≤0.001. <u>Abbreviations:</u> ALT.. Alanin aminotransferase; CK.. Creatine kinase; CSS.. clinical severity score; Ctrl.. Control; h.. hours; hi.. high; *i.p*.. intraperitoneal; LDH.. Lactate dehydrogenase; LPS.. Lipopolysaccharide; Neutrop.. Neurophils; PCI.. peritoneal contamination and infection; PLF.. Peritoneal lavage fluid; *Rag2^−/−^*.. *Rag2*–knock-out.

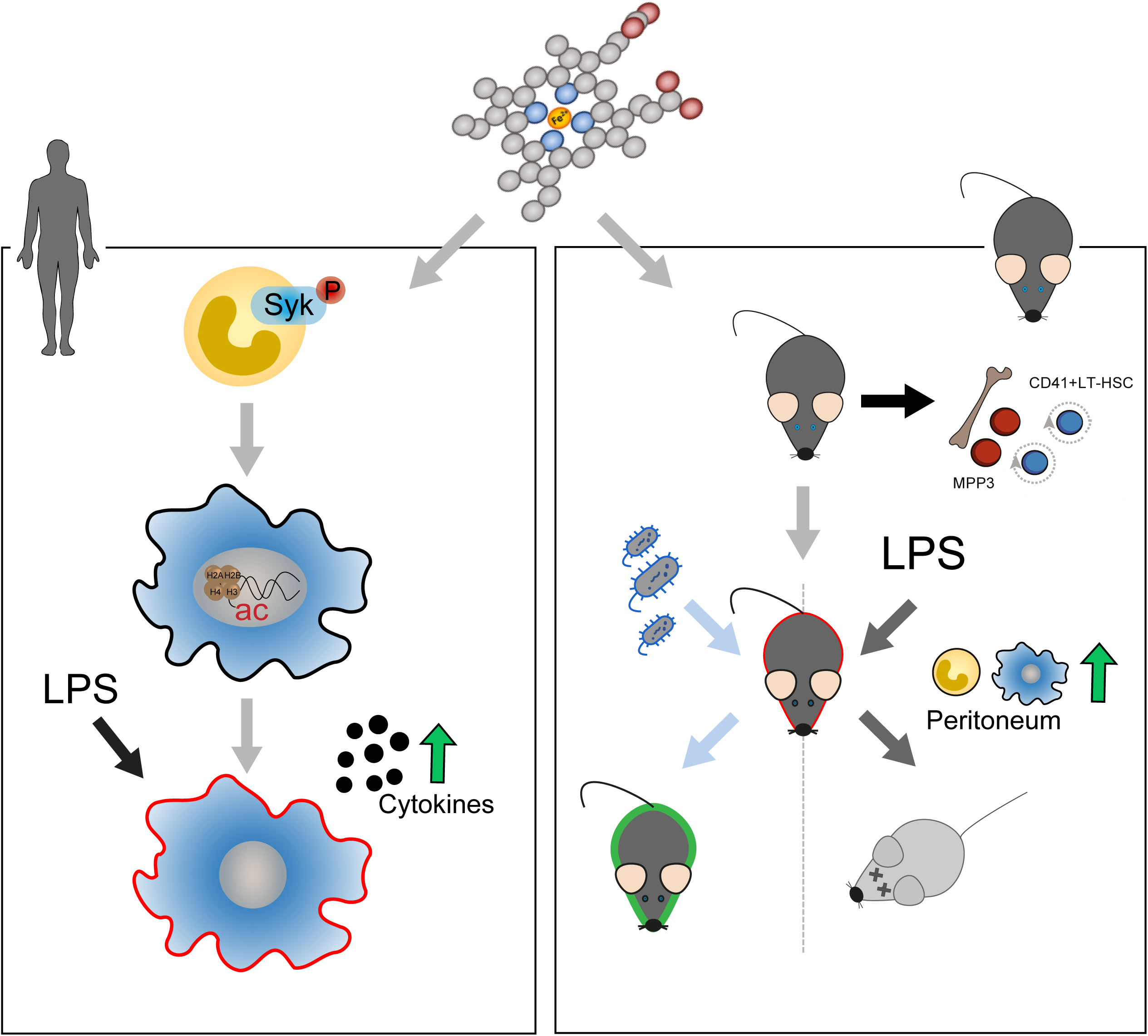

In summary, these data suggest that heme-induced trained immunity promotes a myeloid bias in HPSCs involving self-renewing long term HPSCs. Hence, epigenetic alterations in HSPCs may contribute to the persistence of the heme-induced training effects.

### Heme-induced training improves immunity in bacterial sepsis

We then assessed the effects of heme training *in vivo* using a mouse endotoxemia model (*Fig. 5a*). As a first notion, heme-application alone did not lead to any phenotype in the mice. However, when LPS was injected after 6 days, heme-trained mice displayed increased sickness behavior and increased temperature drops and significantly reduced survival when compared to control mice (*Fig. 5b,c,d*). FACS analysis of immune cell subsets taken from peritoneum 6 hours after LPS application revealed no changes in total cell number, yet, numbers of classical monocytes and numbers of Mφ were significantly higher in heme-trained mice (*Fig. 5e,f*). The deleterious effects of heme training on subsequent LPS-induced systemic inflammation persisted in *Rag2* knock-out mice (*Rag2^−/−^)* with a constitutive deficiency of mature B- and T-cells ^45^ as shown by significantly increased disease severity, significantly decreased temperature and decreased survival of heme-trained *Rag2^−/−^* mice compared to control mice (*Fig. 5g,h,i*). This finding supports that heme-mediated trained immunity is predominantly mediated by effects on innate immune cells. To further assess if heme-induced trained immunity is protective as had been shown for other infectious disease before, we induced a smoldering bacterial sepsis. In contrast to the endotoxemia model, heme-induced trained immunity was protective in polymicrobial sepsis as seen by a faster recovery of weight of heme pre-treated animals as well as a significantly higher temperature and a higher survival rate (*Fig. 5j,k,l*). We coclude that heme-induced trained immunity can be deleterious in acute inflammatory stress but improve the host defense against infections of sutle onset.

## DISCUSSION

Innate immune memory has now been recognized as a fundamental feature of myeloid cells exposed to specific infections, vaccinations or exogenous signals, such as the BCG vaccine, a non-severe *Candida* infection or the fungal cell wall component β-glucan, and certain metabolites ^16–18,46,47^. The relative long-lasting nature of underlying epigenetic and transcriptional programs has led to the concept of “trained immunity”, referring to the capacity of innate immune cells and in particular Mφ to acquire relatively long-lasting immunological memory of past insults ^19,48^. Presumably, this allows innate immune cells to respond to re-challenge by an unrelated pathogen in a more efficient manner both in terms of targeting that pathogen for destruction and limiting the extent of tissue damage associated with this process. The data provided in this manuscript shows that extracellular labile heme, an iron-containing tetrapyrrol alarmin released from red blod cells upon hemolysis acts as a potent inducer of innate immune memory.

When exposed to heme, monocytes accumulate H3K27ac at several hundred regions and regulate the expression of several hundred genes, which can be segregated into two categories. There is a heme-specific inflammatory response and a more general trained immunity response which is shared with β-glucan. It involves lysosome and phagocytosis pathways, related to general monocyte-to-macrophage differentiation, which are central to trained immunity induction. Potentially, this can explain the increased release of cytokines such as TNF upon restimulation. Despite increased TNF secretion in heme-trained Mφ, we could not find a difference in H3K27ac levels at the TNF promoter after heme treatment compared to control cells by ChIP-qPCR which is in line with our finding that the pan-innate immune training signature observed in both heme and β-glucan exposed Mφ involves the upregulation of mechanisms for cytokine release. This is in agreement with recent findings by other authors who used a model of IFNγ-priming ^49^. Increased TNF release was also unrelated to H3K27ac in the TNF promotor itself as IFNγ dependent modifications occur at a region 8 kb upstream of the TNF promotor ^49^. Additionally, it was shown that innate immune memory does not only depend on H3K27ac but also involves histone methylation ^18,50^. This was exemplarily shown for the activating H3K4me3 mark at promotor regions and the H3K4me1 mark at enhancers in β-glucan trained Mφ ^34^, which were not included in our analysis.

Heme-induced training is mediated by a signal transduction pathway involving Syk and JNK phosphorylation and H3K27 acetylation. How heme induces Syk-phosphorylation independently of TLR4 is currently unknown. However, it has been speculated that there is no specific heme receptor involved, as Syk activation can occur via autophosphorylation ^51^ enforced by heme-interaction with phospholipids in the cytoplasmic membrane. To this point, it is also unclear how JNK-signaling leads to histone modifications and which cofactors are involved in this process. Another striking finding is the long-term changes in chromatin accessibility as assessed by single nuclei ATAC sequencing of purified LSK cells. One-time exposure to heme, that did not affect vigor of the animals, resulted in enrichment of chromatin accessibility from genes defining early myeloid progenitors. This suggest that hosts exposed to heme, such as would occur in malaria or other form of hemolysis, prime hematopoeitc stem cells towards adapted myelopoiesis. The heme-dependent transcriptional repressor BACH-1 plays an essential role in the heme induced increase of myeloid cells, since it was shown that heme binding to BACH-1 causes a shift from erythropoiesis towards myelopoiesis in the bone marrow ^52^, similar to observed in this study. Whether the effects observed in our study rely on heme-driven modulation of BACH-1 remains to be established. In mice, innate immune memory induced by β-glucan or BCG was mostly protective ^17,18,23,31^ and is associated with increased myelopoiesis ^25,26^ which is also seen in heme induced innate immune memory. It can be speculated that enhanced cytokine release as a shared feature of both β-glucan and heme-training, explains its protective effect against infection. Mitroulis *et al.* showed that β-glucan training was associated with a specific expansion of myeloid-biased hematopoietic stem cells (HSC)^26^ although HSC do not express the β-glucan receptor Dectin-1. Instead, indirect effects exerted via cytokines seem to be responsible for long-term adaptation of myelopoiesis. Whether this is also the case for heme-training remains to be established.

Trained immunity was initially described after *Candida albicans* and β-glucan application ^17^. Recent data suggests that induction of trained immunity is a common response to different, yet not all, stimuli. While systemic exposure of myeloid cells to β-glucan occurs only rarely, our findings reveal that innate immune traning is a more general component of innate immunity, as hemolysis occurs in very important systemic infectious diseases such as malaria or sepsis ^8^.

## CONCLUSION

Our data show that primary human and murine myeloid cells can undergo heme-training resulting in specific changes of the expression of several hundred genes within 24 hours of exposure. Heme-regulated genes can be segregated into genes encoding cytokines and chemokines versus genes regulating metabolism as well as lysosome and phagocytosis function. The later have also been associated with β-glucan training and may therefore represent a pan-trained immunity signature. Heme imposes histone modifications and chromatin accessibility patterns and relies on signal transduction pathways involving the spleen tyrosine kinase mediated pathway, which is clearly distict from β-glucan-training. This demonstrates that trained immunity can be induced by alarmins, such as heme, which provide long-lasting adaptation of innate immunity to modulate the pathophysiology of a variety of immune-mediated inflammatory diseases.

## METHODS

### Isolation of PBMC and human monocytes

PBMCs were isolated from buffy coats obtained from the Institute of Transfusion Medicine, Jena University Hospital or from freshly donated blood by volunteers (Ethical approval of the Ethics Committee of the University hospital Jena; license reference 4899-07/16). Blood was diluted 1: 7.5 in 1xPBS w/o Ca^2+^ and Mg^2+^ and added on top of Ficoll-Plaque. Layers were separated by centrifugation at 580 g for 30 min and the interphase layer, containing PBMCs, was further processed by washing three times with ice cold PBS. Afterwards the cell pellet was resuspended in warm RPIM and added on top of a hyper-osmotic Percoll solution (48.5 % Percoll, 41.5 % steril H_2_O, 0.16 M filter sterilized NaCl). By centrifugation (15 min; 580 g) layers were separated and interphase layer was washed with ice cold PBS. Cells were resuspended in RPMI culture medium (containing 50 µg/ml gentamicin, 2 mM glutamax and 1 mM pyruvate) and counted. The suspension was adjusted to 10×10^6^ cells per mL with RPMI. Cells were seeded (625,000 cells/cm^2^) on cell culture dishes and incubated for one hour at 37 °C. Cells were washed twice with warm PBS, minimizing contamination by other cells.

### Monocyte Training

Monocyte training was adapted from ^33^. Briefly, after the last washing step, cells were either incubated with culture medium (negative control) or with the respective stimuli for 24 h. Culture medium was supplemented by 10 % pooled human serum. For inhibition experiments, cells were incubated with the respective inhibitor one hour before the stimulation. Inhibitor was present for the whole stimulation time of the first hit. After 24 h cells were washed once with warm PBS and new culture medium with serum was added to the cells. After 48 h medium was exchanged and cells were further incubated for 3 days. On day 6 medium was removed and cells were stimulated with 10 ng/ml LPS in culture medium. A negative control was always included only receiving culture medium. Supernatants were collected 24 h later and stored at −20 °C until determination of cytokines. Cells were either lysed by RIPA buffer and also stored at −20 °C until determination of protein amount or used for RNAseq and lysed in Qiazol.

### Preparation of pooled human serum

Procedures were approved by the Ethics Committee of the University hospital Jena; license reference 4899-07/16. Blood was collected from healthy volunteers in S-Monovette® (Sarstedt). After 15 min resting it was centrifuged at 2,000 g for 10 min. Serum layer was transferred in new tubes and stored at −80 °C. Serum was pooled and centrifuged at 3,500 g for 30 min afterwards it was sterile filtrated and stored at −80 °C until use.

### Isolation of murine bone marrow cells

Bone marrow was isolated of femur and tibia of male mice, 8-12 weeks old. Bones were flushed with RPMI containing 10% FCS and 1% Pen/Strep and stable glutamine. Cells were centrifuged for 7 min at 350xg and 4 °C. Erythrocytes were lysed by resuspending the pellet in red blood cell lysis buffer (150 mM NH_4_Cl; 10 mM KHCO_4_; 130 µM EDTA) for 7 min at room temperature. Reaction was stopped by adding RPMI and cells were again centrifuged. Cells were used directly or stored at −150 °C as cryo stock.

### Differentiation of murine bone marrow derived macrophages

Cells of murine bone marrow were cultured in RPMI containing 10% FCS and 1% Pen/Strep. Differentiation was induced by adding 20 ng/ml macrophage colony stimulating factor (M-CSF; Peprotech). One week later cells were harvested and seeded on well plates and incubated for one day with GM-CSF (20 ng/mL; Peprotech). Training experiments were performed with these cells according to the Monocyte training protocol. During this time 0.2 ng/mL GM-CSF was always added to the medium.

### Pharmacological Approaches

Epigenetic modifications were inhibited as previously used ^53^. Following enzymes were blocked Histone acetyltransferases by epigallocatechin-3-gallate (EGCG; 15 µM; Sigma-Aldrich), histone methyltransferases by 5′-deoxy-5′-methylthioadenosine (MTA; 1mM; Sigma-Aldrich) and histone demethylasis by pargyline (3 µM; Sigma-Aldrich). Pathway inhibition was realized by using specific inhibitors. Accordingly, R406 (1 µM; InvivoGen) was used for inhibiting Spleen tyrosine kinase activation. JNK pathway was blocked by SP600125 (10 µM; Invivogen). Rapamycin (10 nM; InvivoGen) was used for blocking mTOR mediated signal transduction. Reactive oxygen species (ROS) were scavenged by N-acetylcystein (10 mM; Fluka). ROS measurments were performed by incubating the cells with H2DCFDA and flow cytometry analysis. Ultrapure LPS from *E.coli* O55:B5 was purchased from Inviogen. ATP was purchased from InvivoGen. Stimulation with heme (Sigma-Aldrich) and PPIX was, if not otherwise stated, performed with 50 µM. For β-glucan training cells were stimulated with 1 µg/mL β-glucan.

### Tetrapyrrole preparation

Stock solution of Heme (Sigma-Aldrich for cell culture experiments and Frontier Scientific for animal experiments) and Protoporphyrin 9 (PPIX; Frontier Scientific) was prepared as described in ^2^. Briefly, it was dissolved in 0.2 M NaOH until it was completely dissolved. Afterwards, 0.2 M HCl was added slowly and solution was adjusted to pH7.4. Afterwards, the solution was centrifuged at 4,000 rpm; 15 min 4 °C and sterile filtered with a 70 µM cell strainer. Concentration of the stock was determined by measuring OD_405_ of a 1:1,000 dilution in DMSO.

### Animal Experiments

Experimental procedures were ethically reviewed and approved by the Ethics Committee of the Instituto Gulbenkian de Ciência (license reference: A009/2011, by Direção Geral de Alimentação e Veterinária (license reference: 0420/000/000/2012) or by the regional animal welfare committee (Registration number: 02-044/16, Thuringian State Office for Consumer Protection and Food Safety). All experiments conducted on animals followed the Portuguese (Decreto-Lei nº 113/2013) or the German legislation on protection of animals and the European (Directive 2010/63/EU) legislations, concerning housing, husbandry and animal welfare. C57Bl/6 mice were bred and maintained under specific pathogen free conditions at the Instituto Gulbenkian de Ciência or at the Jena University Hospital (12 h day/night; fed *ad libidum*).

*Tlr4*^−/−^ (obtained originally from Shizuo Akira, Osaka University, Japan) and *Rag2*^−/−^ mice were breed at the Instituto Gulbenkian de Ciência, Oeiras, Portugal. Mice received 2 mg/kg heme (Frontier Scientific) or vehicle (NaOH; pH 7.4) intraperitoneal (*i.p*.). For survival experiments and flow cytometry analysis of peritoneal lavage fluid cells mice received 15 mg/kg LPS (*E.coli* O55:B5; purified by phenol extraction; Sigma-Aldrich) one week after heme treatment. Temperature, weight and clinical severity score ^10^ was monitored for one week.

### Cytokine measurements

Cytokines were determined in supernatants of cells treated 24 h with LPS. TNF, as regular read-out, was determined using ELISA kits following the manufacturer’s instructions. In addition, cytokines in supernatants and in mouse plasma were determined using the LegendPlex™ Human Inflammation Panel (13-plex) or the LegendPlex™ Mouse Inflammation Panel (13-plex) respectively according to manufacturer instructions. Cytokine values were normalized to protein values of respective wells. Only samples which clearly showed training in TNF ELISA after normalization to protein were taken for the LegendPlex™ analysis.

### Serology

Serological parameters were determined by SYNLAB.vet GmbH, Leipzig, Germany.

### Western Blot

Proteins concentrations were determined by Bicinchoninic acid (BCA) assay (Serva), performed according to manufacturer instructions. Samples were separated in a 10 % acrylamide gel and blotted onto a PVDF membrane (Roth, Germany). Unspecific binding was blocked with 5 % BSA in TBS-Tween for one 1 h. The following primary antibodies were used overnight at 4 °C: p-Syk (1:200, rabbit, Cell signaling technologies (CST), AB_2197222), Syk (1:1,000, rabbit, CST, 4D10), p-JNK (1:1,000, rabbit, CST, AB_10831195), JNK (1:1,000, rabbit, CST, AB_10694056), p-S6 (1:1,000, rabbit, CST, AB_331679), S6 (1:1,000 rabbit, CST, AB_331355), and GAPDH (1:20,000 rabbit, CST, AB_561053) and proteins were detected by subsequent incubation in peroxidase conjugated secondary antibodies (2 h; room temperature) and developed with SERVALight Eos CL HRP WB Substrat-Kit (Serva). ImageJ was used for quantification ^54^. For Western blot analyses of H3K27ac, cells were lysed in RIPA buffer supplemented with inhibitors (Phenylmethylsulfonyl fluoride (PMSF) (Carl Roth), Protease inhibitor cocktail (PIC) complete ULTRA tablets mini (Roche), Nicotinamind (NAM) (Sigma) and Trichostatin A (TSA) (Sigma) and sonicated by Sonifier W-250D (Branson) (25% amplitude, 30 sec (5 sec on, 5 sec off)). Samples were separated by a denaturating 15% SDS polyacrylamide gel with subsequent blotting onto a PVDF membrane. As blocking reagent 5 % skim milk powder (Saliter) in PBS-T was used for 1 h at room temperature (RT). Blots were incubated over night at 4°C with the following primary antibodies diluted in 2 % skim milk powder/PBS-T: H3K27ac (1:1,000, rabbit, Active Motif, 39133) and H3 (1:10,000, rabbit, ab1791). Peroxidase conjugated secondary antibodies (1:7,500, goat, Thermo Fisher, 31460) in 2 % skim milk powder/PBS-T were applied for 1 h at room temperature. For chemiluminescent reaction Cyanagen Westar Nova 2.0 (XLS071,0250) or Westar Eta C Ultra 2.0 (XLS075,0100) was used and the reaction was detected by Fusion Solo S (Vilber). Quantification of the blots was perfomred using BIO-1D advanced (Vilber) Software.

### RNA-Sequencing

Total RNA was extracted from cells using the Qiagen RNeasy RNA extraction kit (Qiagen, Netherlands), using on-column DNAseI treatment. Illumina library preparation was done using the RNA HyperPrep Kit with RiboErase (KAPA Biosystems). Sequencing was performed using the Illumina NextSeq500 machine 75bp 400M reads kit.

### Chromatin Immunoprecipitation and Sequencing (ChIP-seq)

Purified cells were fixed with 1% formaldehyde (Sigma) at a concentration of approximately 10 million cells/ml. Fixed cell preparations were sonicated using a Diagenode Bioruptor Pico 5 minutes (30 seconds on; 30 seconds off). 67 µL of chromatin (1 million cells) was incubated with 229 µL dilution buffer, 3 µL protease inhibitor cocktail 1µg of H3K27ac antibody (Diagenode) and incubated overnight at 4 ºC with rotation. Protein A/G magnetic beads were washed in dilution buffer with 0.15% SDS and 0.1% BSA, added to the chromatin/antibody mix and rotated for 60 minutes at 4 °C. Beads were washed with 400 µL buffer for 5 minutes at 4 °C with five rounds of washes. After washing, chromatin was eluted using elution buffer for 20 minutes. Supernatant was collected, 8 µL 5M NaCl, 3µL proteinase K were added and samples were incubated for 4 hours at 65 °C. Finally, samples were purified using Qiagen; Qiaquick MinElute PCR purification Kit and eluted in 20 µL EB. Illumina library preparation was done using the Kapa Hyper Prep Kit. For end repair and A-tailing double stranded DNA was incubated with end repair and A-tailing buffer and enzyme and incubated first for 30 minutes at 20 °C and then for 30 minutes at 65 °C. Subsequently adapters were ligated by adding 30 µl ligation buffer, 10 Kapa l DNA ligase, 5 µL diluted adaptor in a total volume of 110 µL and incubated for 15 minutes at 15 °C. Post-ligation cleanup was performed using Agencourt AMPure XP reagent and products were eluted in 20 µl elution buffer. Libraries were amplified by adding 25 µL 2x KAPA HiFi Hotstart ReadyMix and 5 µL 10x Library Amplification Primer Mix and PCR, 10 cycles. Samples were purified using the QIAquick MinElute PCR purification kit and 300 bp fragments selected using E-gel. Correct size selection was confirmed by BioAnalyzer analysis. Sequencing was performed using the Illumina NextSeq500 machine 75 bp 400M reads kit. Detailed protocols can be found on the Blueprint website (http://www.blueprint-epigenome.eu/UserFiles/file/Protocols/Histone_ChIP_May2013.pdf). Enrichment was calculated as percentage of total chromatin input and fold change was calculated by normalizing treated samples to untreated controls.

### Nuclei isolation and single-nuclei ATAC library construction

LSK cells (Lin^−^cKit^+^Sca1^+^) were sorted from untreated (n = 2), and Heme treated (n = 2) mice. Isolation was performed following the recommended guidelines from 10x Genomics for low cell input with some slight modifications. Briefly, after thawing LSK cells at 37°C for 2 minutes, they were transferred to a 2 mL microcentrifuge tube and added 1 mL of PBS + 0.04% BSA (resuspension buffer). Cells were immediately centrifuged at 300 g for 5 min at 4°C. The supernatant was removed carefully, then 100µL chilled resuspension buffer were added and transferred the cell suspension to a 0.2 mL tube. Cells were again centrifuged (300 g for 5 min at 4°C), followed by removal of 90 µL supernatant. 50µL of chilled lysis buffer (10 mM Tris-HCl (pH 7.4), 10 mM NaCl, 3 mM MgCl_2_, 0.1% Tween-20, 0.1% Nonidet P40 Substitute, 0.01% digitonin and 1% BSA) was added, gently pipette-mixed and incubated on ice for 3 min. Nuclei were centrifuged at 500 g for 5 min at 4°C and 45 µL of supernatant removed without disrupting the nuclei pellet. Nuclei were resuspended in 45 µL chilled Diluted Nuclei Buffer (10x Genomics; 2000153) without mixing. Followed by another centrifuging step, the supernatant was removed completely, and nuclei were resuspended in 12 µL of Diluted Nuclei Buffer. 2 µL were taken to calculate nuclei concentration using a Countess II FL Automated Cell Counter. The nuclei concentration was between 6,000 to 8,000 nuclei per µL.

For a detailed protocol for library preparation, refer to the Chromium Single Cell ATAC v1 User Guide. Upfront tagmentation of ~35,000 nuclei was performed for 1 h at 37°C in a C1000 Touch thermal cycler with 96-Deep Well Reaction Module (Bio-Rad, 1851197). Tagmented nuclei were mixed the master mix and loaded onto a Chromium Chip with Gel beads and partitioning oil on different wells. The chip was placed into a Chromium Single Cell Controller instrument (10x Genomics). The samples were processed on the same day to avoid potential technical bias. Resulting single-nuclei GEMs were collected and PCR amplification was performed in a C1000 Touch Thermal cycler as follows: 72 °C for 5 min, 98 °C for 30 s, cycled 12×: 98 °C for 10 s, 59 °C for 30 s and 72 °C for 1 min. GEMs were broken, and products were cleaned up using Dynabeads and SPRIselect reagent (Beckman Coulter; B23318). Indexed libraries were obtained after mixing the barcoded amplicons with the Sample Index PCR Mix and amplified following the next PCR conditions: 98 °C for 45 s, cycled 11x: 98 °C for 20 s, 67 °C for 30 s, 72 °C for 20 s, and 72 °C for 1 min. The sequencing libraries were clean-up with SPRIselect reagent, and the nucleosome pattern was assessed on a 2100 Bioanalyzer Instrument (Agilent, G2943CA) using a High Sensitivity DNA chip (Agilent, 50674626). DNA concentration was additionally quantified by Qubit 1x dsDNA HS Assay Kit (ThermoFisher, Q33230). Libraries were loaded on a NextSeq 500 (Illumina, SY-415-1001) using the NextSeq High Output Kit (Illumina 75 cycles, 20024911) and sequenced following the next protocol: read 1, 38 bp; i7 index read, 8 cycles; i5 index read, 16 bp; and read 2, 38 bp.

### ChIP-qPCR

Isolated cells were thawed and subsequently washed two times with PBS. 1×10^6^ to 5×10^6^ cells were fixed with 1% formaldehyde for 10 min. Subsequent chromatin preparation steps were performed according to the “High Sensitivity Chromatin Preparation” Kit (Active Motif, 53046). Samples were sonicated two times for 10 minutes (30 seconds on, 30 seconds off) by Diagenode Bioruptor Pico. Protein A and Protein G coupled magnetic beads (mixture 1:1, Dynabeads, Thermo Fisher, 10002D and 10004D) were pretreated with BSA (NEB, B9001) and DNA from herring sperm (Sigma, D7290), rotating for 1 h at 4°C. Then beads were incubated with H3K27ac antibody (1 µg per IP, rabbit, Active Motif, 39133) or normal rabbit IgG (1 µg per IP, rabbit, CST, 2729) for 4 h at 4°C and rotating. To each sonicated sample 300 µL ChIP Buffer (Active Motif) supplemented with PMSF (Carl Roth), PIC (Roche), NAM (Sigma) and TSA (Sigma) was added. Then, samples were incubated with H3K27ac-coupled magnetic beads and normal rabbit IgG-coupled magnetic beads overnight at 4°C and rotating. The next day, beads were washed with wash buffer 1 (20 mM Tris-HCl pH 8, 150 mM NaCl, 2 mM EDTA, 1 % Triton X-100, 0.1 % SDS), wash buffer 2 (20 mM Tris-HCl pH 8, 500 mM NaCl, 2 mM EDTA, 1 % Triton X-100, 0.1 % SDS), wash buffer 3 (10 mM Tris-HCl pH 8, 250 mM LiCl, 1 mM EDTA, 1 % NP40, 1 % Sodium deoxycholate) and two times with wash buffer 4 (10 mM Tris-HCl, pH 8, 1 mM EDTA). For elution, beads were incubated two times with 40 µl elution buffer (100 mM NaHCO3, 1 % SDS) for 5 minutes rotating on an Intelli-Mixer (neoLab). After elution, samples were incubated with 1 µL RNase (1µg/µL) (Sigma) at 37°C for 1 h and subsequently incubated with 2 µL Proteinase K (20 mg/ml) (Thermo, EO0491) at 56°C for 1 h. For decrosslinking, 5 M NaCl was added to a final concentration of 250 mM and samples were incubated at 65°C overnight. DNA was purified using the ChIP DNA Clean & Concentrator Kit (Zymo Research, 5205) and stored at −20 °C for qPCR analysis. qPCR was performed on a StepOne Plus system (Applied Biosystems) using primers with a concentration of 100 nM (*IL8*) and the PowerUp™ SYBR™ Green Master Mix (Thermo, A25780). qPCR measurements were performed as triplicates and amplification procedure was as follows: step 1: 95°C for 10 minutes, step 2: 95°C for 15 seconds, step 3: 60°C for 1 minute, step 4: 95°C for 15 seconds, step 5: 60°C for 1 minute, step 6: 60-95°C ramp for +0.3°C, step 7: 95°C for 15 seconds, step 2 and 3: 40 cycles. Samples were normalized to the percentage of Input DNA by using %Input = 100*2^(Ct_INPUT_-Ct_IP_). % Input of each treated sample was then normalized to the untreated control. Following Primer Sequences were used: *IL8*_ChIP_fw3: GCAGAGCTGTGCCTGTTGAT *IL8*_ChIP_re3: CCTTGGCAAAACTGCACCTG.

### Flow cytometry

#### Hematopoietic stem and progenitor cells

Cell analysis was performed on a FACS Canto II flow cytometer (BD, Heidelberg, Germany) using FACSDiva 6.1.3 software (BD). Bone marrow (BM) cells were harvested from femurs by repeatedly flushing with PBS supplemented with 5% FBS (Life Technologies). The analysis of BM cellularity was performed with MACSQuant (Miltenyi Biotec). Antibodies to Sca1 (clone E13-161.7), cKit (clone 2B8), anti-CD135 (Flt3, clone A2F10), anti-CD48 (clone HM48-1), anti-CD41 (clone MWReg30), anti-CD150 (clone TC15-12F12.2), anti-CD16/CD32 (clone 93), anti-CD34 (clone RAM34) as well as a lineage cocktail (Lin; anti-CD3e, anti-Ly6G/Ly6C, anti-CD11b, anti-B220 and anti-Ter-119) were used for cell surface phenotype analysis. Data analysis was performed using FlowJo™ (Tree Star) software.

#### LSK sorting for single cell ATAC-sequencing

Cell sorting was performed using a FACS Aria III cell sorter (BD, Heidelberg, Germany) ^55–57^. Cell purity was above 95%. Single cell suspensions from bone marrow were prepared. Enrichment of Lin^−^ cells was performed prior to cell sorting by negative magnetic selection using MidiMACS Separator (Miltenyi Biotec). Cells were incubated with a biotin mouse lineage panel (Biotin-Conjugated Mouse Lineage Panel, BD PharMingen) and subsequently with anti-Biotin MicroBeads (Miltenyi Biotec). For cell surface staining a lineage cocktail (Lin), including anti-CD3e (clone 145-2C11), anti-CD11b (clone M1/70), anti-Gr1 (clone RB6-8C5), anti-B220 (clone RA3-6B2), and anti-TER119 (clone TER-119), anti-CD117 (clone: 2B8), anti-Sca1 (clone E13-161.7). LSK cells were sorted as Lin^−^CD117^+^Sca1^+^.

#### Peritoneal cells

Analysis of cells from peritoneal lavage fluid was performed on an LSR Fortessa X20 flow cytometer (BD, Heidelberg, Germany) using FACSDiva 8.0.1 software (BD). Peritoneal lavage fluid was collected from the peritoneal cavity of mice pretreated with vehicle or heme and treated with LPS for 6h. Peritoneal lavage fluid cells were counted using a Neubauer hemocytometer and stained with an antiboy mix for cell surface phenotype analysis. The antibodies anti-CD11b (clone M1/70), anti F4/80 (clone BM8), anti-Ly6G (clone 1A8), anti-Ly6C (clone HK1.4), anti MMR (clone C068C2) and anti-MHCII (clone M5/114.15.2) were used. Cell doublets and debris were excluded using FSC/SSC gates and viable cells were discriminated from dead cells using Live/dead fixable yellow dye (Life Technologies).

### Data and Software Availability

Raw data files for the RNA, ChIP and single cell ATAC sequencing and analysis have been deposited as a Series in the NCBI Gene Expression Omnibus under accession number: <u>GSE111003</u>. The link to GEO sessions is: https://www.ncbi.nlm.nih.gov/geo/query/acc.cgi?acc=GSE111003.

### Quantification and Statistical Analysis

Survival Plots were analyzed using the Log-rank test. One-Way ANOVA with Fisher’s LSD was used to analyse more than two groups. Two-Way-ANOVA with Fisher’s LSD was used when analyzing more than one group and time point. Student t-Test was used for pair-wise comparisons. FACS data analysis was performed using FlowJo (Tree Star) software.

#### RNA-seq analysis

To infer gene expression levels, RNA-seq reads were aligned to the Ensembl v68 human transcriptome using Bowtie 1. Quantification of gene expression was performed using MMSEQ. Differential expression was determined using DEseq. Differentially expressed genes were determined using pair-wise comparison between treatments, with a fold change >2.5, adjusted p value of <0.05, and RPKM ≥5.

#### ChIP-seq analysis

Sequencing reads were aligned to human genome assembly hg19 (NCBI version 37) using bwa. Duplicate and low-quality reads were removed after alignment using Samtools and Bamtools. For peak calling the BAM files were first filtered to remove the reads with mapping quality less than 15, followed by fragment size modeling. MACS2 was used to call the peaks. H3K27ac peaks were called using the default (narrow) setting. Reads/peak tables were merged using Bedtools. Data were normalized using the R package DESeq1 and then pair-wise comparisons were performed (fold change >1.5, p value <0.05 and mean reads/peak ≥20 in any condition) to determine the differential peaks.

#### snATAC-seq data pre-processing and analysis

BCL files were demultiplexed by Cell Ranger

ATAC (v.1.1.0) using ‘cellranger-atac mkfastq’. FASTQ files were aligned to the mm10 reference genome using ‘cellranger-atac count’ with default parameters. This pipeline also marks duplicated reads and identifies transposase cut sites. Cells were filtered by TSS enrichment and unique fragments, as previously described ^58^. This approach allows the detection of TSS sites without having a defined peak set. Cells were filtered, keeping those with a TSS enrichment of at least five and more than 1,000 unique fragments. Since the peak calling was performed separately, peaks do not overlap perfectly and might be treated as entirely different features. Using ‘GenomicRanges’ we find overlaps of the chromosome coordinates among samples. The obtained count matrix was used to generate a union peak set, first binarizing the top 20,000 most accessible sites followed by term frequency-inverse document frequency (TF-IDF) transformation. We then used singular value decomposition (SVD) implementation and created a Seurat ^59^ (v.3.1) object for non-linear dimension reduction. LSI scores were then used for clustering, and visual representation of the clusters was performed using a UMAP projection. The contribution of each cell type per cluster was plotted using ggplot2 in R. To annotate the cell clusters, we used the union peak set and looked for enrichment of epigenomic signatures using CisTopic (v.0.2.2) ^60^. Bulk ATAC profiles of purified cell types from the mouse immune system were used as a reference. Additionally, each accessible region was linked to the close gene base on CHIPseeker annotations, and the probability of each region to a defined set of marker genes was depicted as an AUC score and plotted with CisTopic. Next, we aggregate cells from each condition to create an ensemble track for peak calling and visualization using MACS2. The obtained BedGraph files were compressed to bigWig files using bedGraphToBigWig and explored with WashU Epigenome browser.

## CONTACT FOR REAGENT AND RESOURCE SHARING

Further information and requests for reagents and resources can be directed to, and will be fulfilled by, the Lead Contact, Sebastian Weis.

## ACKNOWLEDGMENTS

We thank Dr. David L. Williams, Ph.D. from the Quillen College of Medicine East Tennessee State University, USA for providing the β-glucan used in this study.

## AUTHOR CONTRIBUTIONS

**EJ** performed *in vitro* experiments, contributed to the study design, performed animal experiments and wrote the manuscript. **BN** performed RNAseq, ChIPSeq and the bioinformatics analysis. **CR** performed scATACseq and bioinformatics analysis. **IK, LK, AE** and **TG** analyzed and interpreted bone marrow progenitor data; **RM** performed animal experiments and FACS analysis. **TC** designed and interpreted mouse progenitor studies and edited the manuscript. **JG** performed western blot analysis. **MaB** performed western blot analysis. **FR** performed mice *in vitro* experiments. **PB** performed ChIP-qPCR and Histone western blots. **MG, HGS, LABJ, TC, MPS, MGN, MB** contributed critically to study design, performed and/or contributed to experiments. **SW** formulated the original hypothesis, drove the study design, designed experiments and wrote the manuscript together with EJ. All authors read and approved the manuscript.

## ABBREVIATIONS

HO-1/*HMOX1*: heme oxygenase 1
IL: Interleukin
LPS: lipopolysaccharide
Mφ: macrophage
PPIX: protoporphyrin IX
ROS: reactive oxygen species
TNF: tumor necrosis factor
TLR: toll like receptor

