## Supplementary Figure Legends for "Heme induces innate immune memory"

**Supplementary Figure 1** (Related to *Figure 1*)**: Heme induces trained immunity in human macrophages**

**(a)** Serum and dose dependent heme-training of human Mφ. Serum concentrations were only modified for the 1^st^ hit. During resting period 10 % serum was present in the culture medium. TNF release after LPS re-stimulation of vehicle or heme-trained Mφ. n=4 independent experiments. **(b)** Cytokine release after LPS re-stimulation of vehicle or heme-trained Mφ. n=16 individual donors. TNF release after LPS re-stimulation from **(c)** vehicle or LPS trained Mφ. n=10 independent experiment; **(d)** vehicle or ATP-trained Mφ. n=2 independent experiment, **(e)** or from heme (50 µM) re-stimulation from vehicle or heme-trained Mφ, n=2 independent experiments, mean±SD, **(f)** vehicle, heme or PPIX trained Mφ, n=4 independent experiments, **(g)** vehicle or heme-trained Mφ with or without ROS- scavenger NAC, n=3 independent experiment. **(h)** ROS production after LPS re-stimulation of vehicle or heme-trained Mφ, n=4 independent experiment **(i)** ROS production of heme with or without NAC (10 mM) treated Mφ 4 h after treatment, n=2 independent experiment. mean±SD**. (j)** Experimental setup*.* **(k)** Representative HO-1 Western blot of vehicle and heme-trained BMDM 24 h after LPS application. **(l)** TNF production of heme-trained *Tlr4^-/-^* mice that received the TLR1/2 agonist Pam3CSK as 2^nd^ hit. n=3 mice. Students t-test; mean±SD.

All ELISA data were normalized to total protein and fold change was calculated to the vehicle control 24 h after LPS treatment. N= 2-3 donors. Data is mean±SEM unless otherwise stated. Student’s t-test or One-way ANOVA with Fisher’s LSD. *p≤0.05; **p≤0.01; ***p≤0.001.

Abbreviations: ATP.. Adenosine triphosphate; Ctrl.. Control; LPS.. Lipopolysaccharide; NAC.. N- acetylcysteine; n.d… not detectable; n.s.. non-significant; PPIX.. Protoporphyrin IX; TNF.. Tumor necrosis factor.

**Supplementary Figure 2** (Related to *Figure 1 and Figure 2*)**: Heme training induces epigenetic modifications in human myeloid cells and heme induced macrophage polarization.**

**(a)** Dynamic regulation of H3K27ac marks. (**b)** log2 tag acetylation intensity of β-glucan trained myeloid cells. (**c,d)** H3k27ac z-score of (c) heme or (d) β-glucan specifically regulated marks. (**e)** Most dynamic genes with β-glucan. (**f)** H3K27ac location relative to TSS. (**g)** GSEA of H3K27ac peaks increased in heme-trained Mφ, Top 10. (**h)** GSEA of H3K27ac peaks decreased in heme-trained Mφ. (**h-j)** Transcriptional analysis of polarization markers for (**h**) M1-like Mφ; (**i**) M2-like Mφ and (**j**) M-heme-like Mφ. All analysis from n=5 individual donors.

Abbreviations: β-Glu, β-glucan; Enh.. Enhancer; h.. hours; H3K27ac.. Histone 3 Lysine 27 acetylation; Prom.. Promotor; TSS.. transcriptional start side.

**Supplementary Figure 3** (Related to *Figure 2*)**: Donor specific heme and β-glucan induced transcriptional modifications**

**(a)** *FOSL1* RNA expression of 5 individual donors after heme or β-glucan treatment and without treatment over time. Extracted data from RNAseq analysis as an examplary gene for donor specific regulation. **(b)** Exemplary motif regulation of heme-specific; β-glucan specific and commonly regulated motifs over time. Data shown as mean (grey line) and standard deviation (shaded area) from 5 independent donors. **(c)** Examplary mRNA expression of heme/iron regulated genes. Extracted data from RNAseq analysis. Data shown as mean (grey line) and standard deviation (shaded area) from 5 independent donors. **(d,e)** Densitometry of time- dependent (d) Syk phosphorylation and (e) JNK phosphorylation n=3 individual donors, **(f)** Western blot and densitometry of p-S6 24 h after heme training of human monocytes with or without Syk- Inhibitor R406. Phosphorylation densitometry signals were normalized to the respective pan-protein and to the housekeeper.

**Supplementary Figure 4** (Related to *Figure 3 and Figure 4*)**:**

**(a)** Experimental setup **(b)** Gating strategy of LSK, CD41^+^-LT-HSC and MPPs. **(c)** Total amount per femur of neutrophils and relative amounts of MEP; CMP; GMP of vehicle or heme-trained mice at day 7. n=6 animals per group derived from 2 independent experiments. **(g)** Total or relative amount per femur of HSPCs of vehicle or heme-trained mice at day 28. n=4 animals per group.

All densitometries were analyzed by One-way ANOVA with Fisher’s LSD; mean ± SEM. All HSPC analysis were analyzed by Student’s t-test. *p≤0.05; ** p≤0.01.

Abbreviations: CMP.. common myeloid progenitor; d.. days; GMP.. Granulocyte-macrophage progenitor; HSC.. hematopoietic stem cell; JNK-.. c-Jun- N-terminal kinases; LSK..Lin^-^Sca1^+^cKit^+^; LT.. long term; MPP.. multipotent progenitor cells; n.s..non-significant; Syk.. Spleen tyrosine kinase; S6..ribosomal protein S6; ST.. short term.

**Supplementary Figure 5** (Related to *Figure 4*)**: Gene scores of single-nuclei ATAC-sequencing from LSK cells.** Projection of gene accessibility scores of known markers of myeloid and megakaryocyte-erythrocyte progenitors from sorted LSK cells. Below each UMAP projection, a genome track window of ~200kb shows the chromatin accessibility of each gene.

**Supplementary Figure 6** (Related to *Figure 4*)**: Epigenomic signatures of single-nuclei ATAC-sequencing from LSK cells.** Projection of the enrichment of epigenomic signatures (AUC scores) for bulk ATAC-seq profiles of early progenitors from the ImmGen Database onto UMAP plots at 7 and 28d.
