## Supplementary figures and images for "Heme induces innate immune memory"

### Suppl.Fig.1

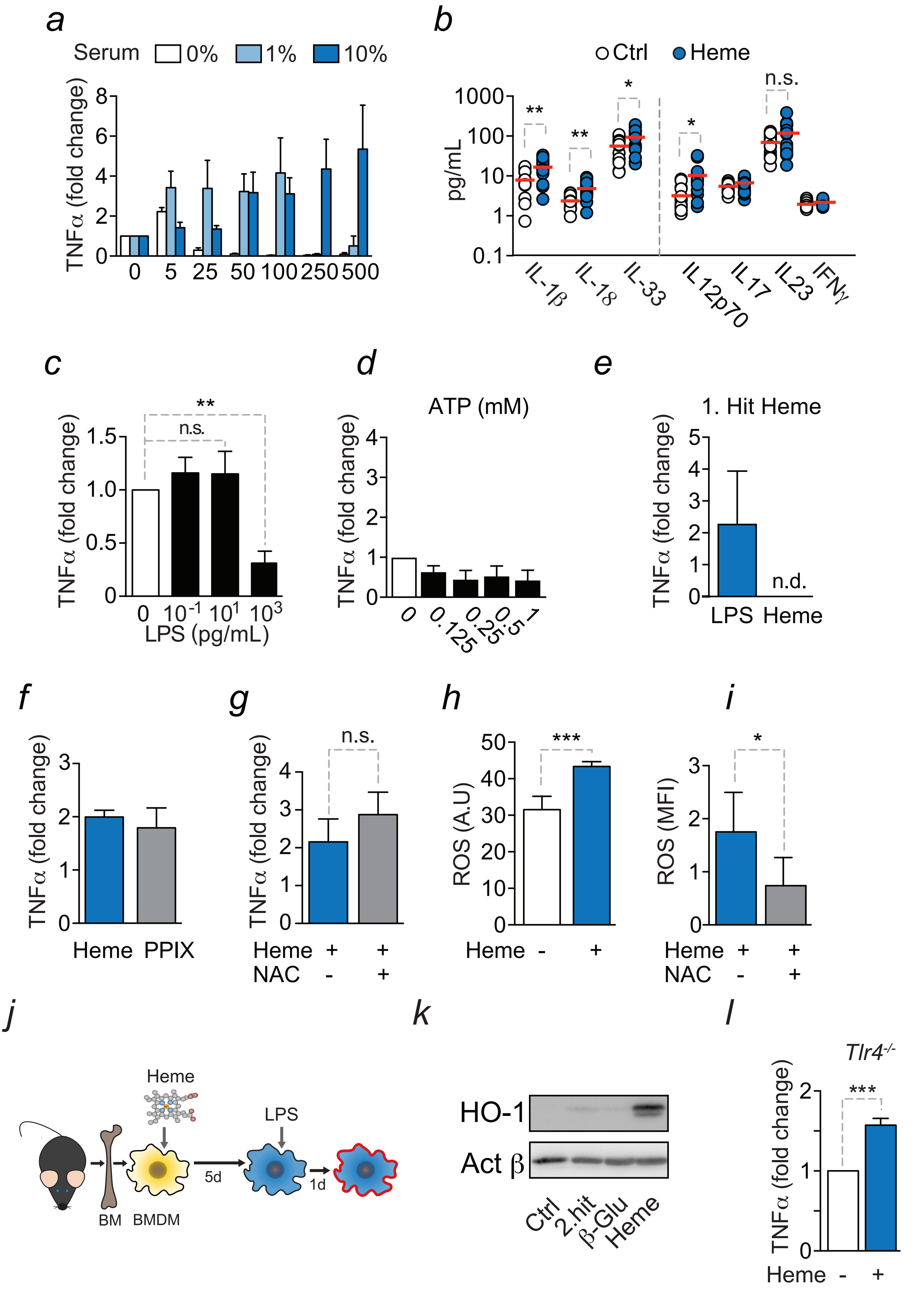

### Suppl.Fig.2

Jentho *et al.* Suppl.Fig. 2

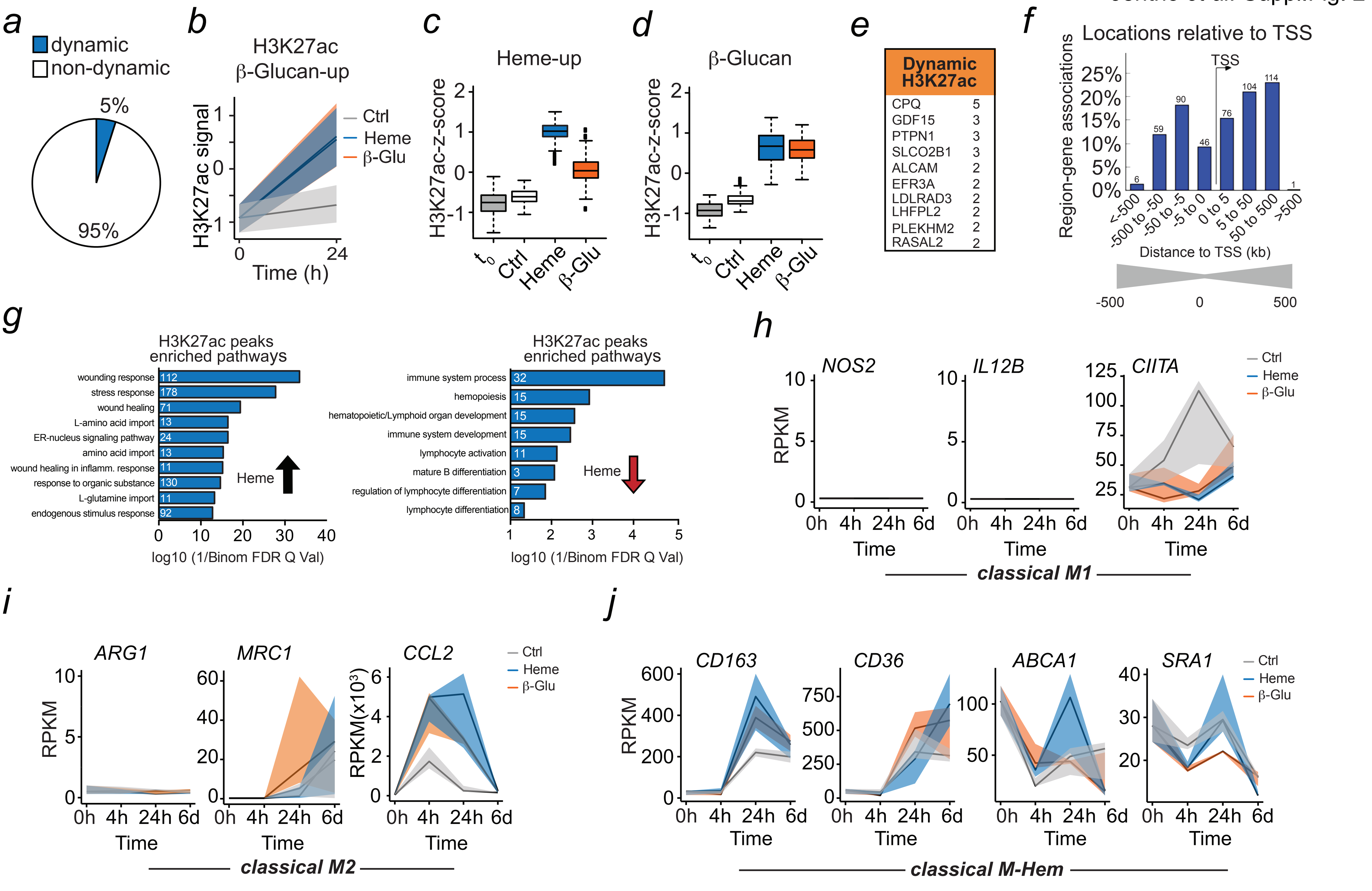

### Suppl.Fig.3

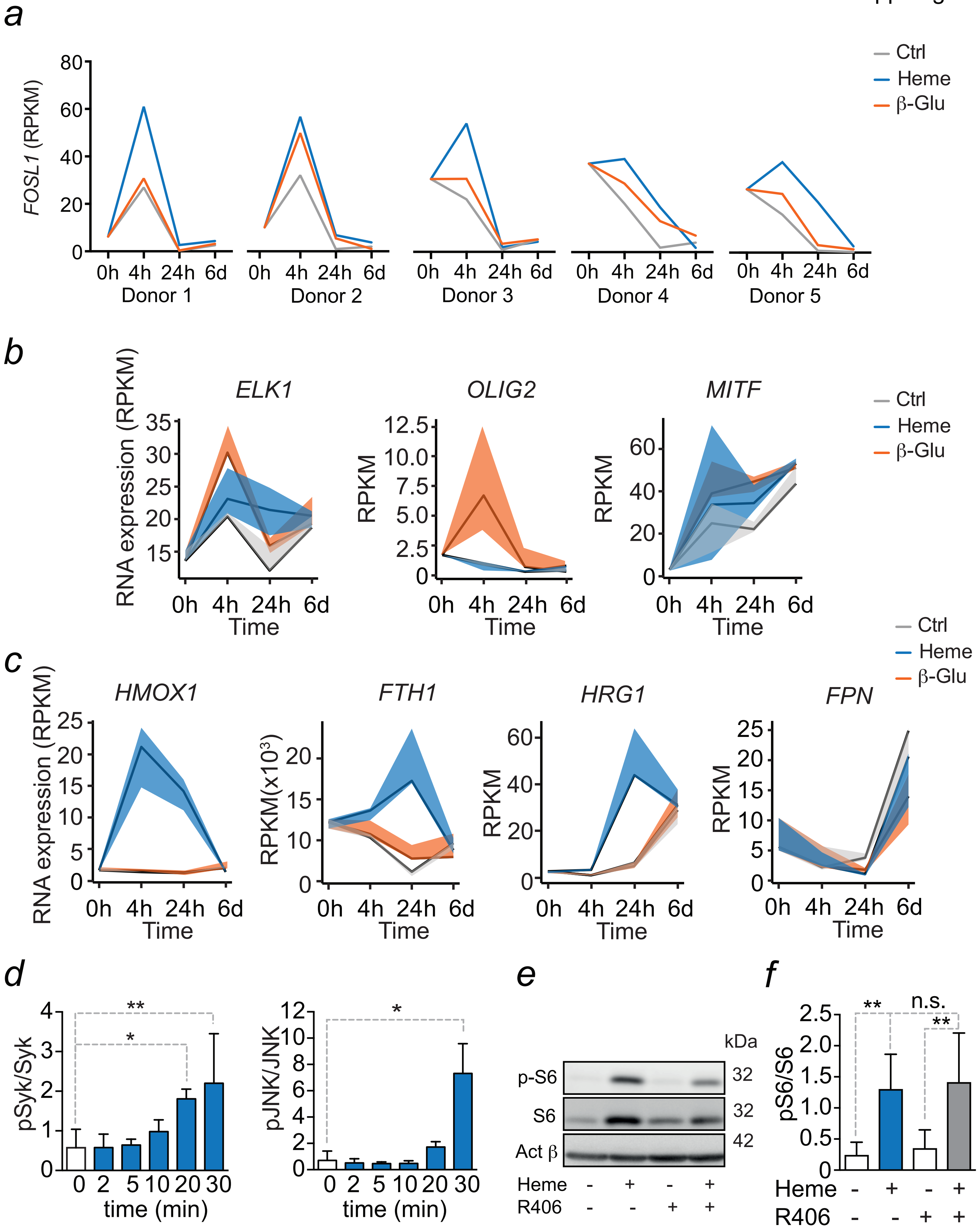

### Suppl.Fig.4

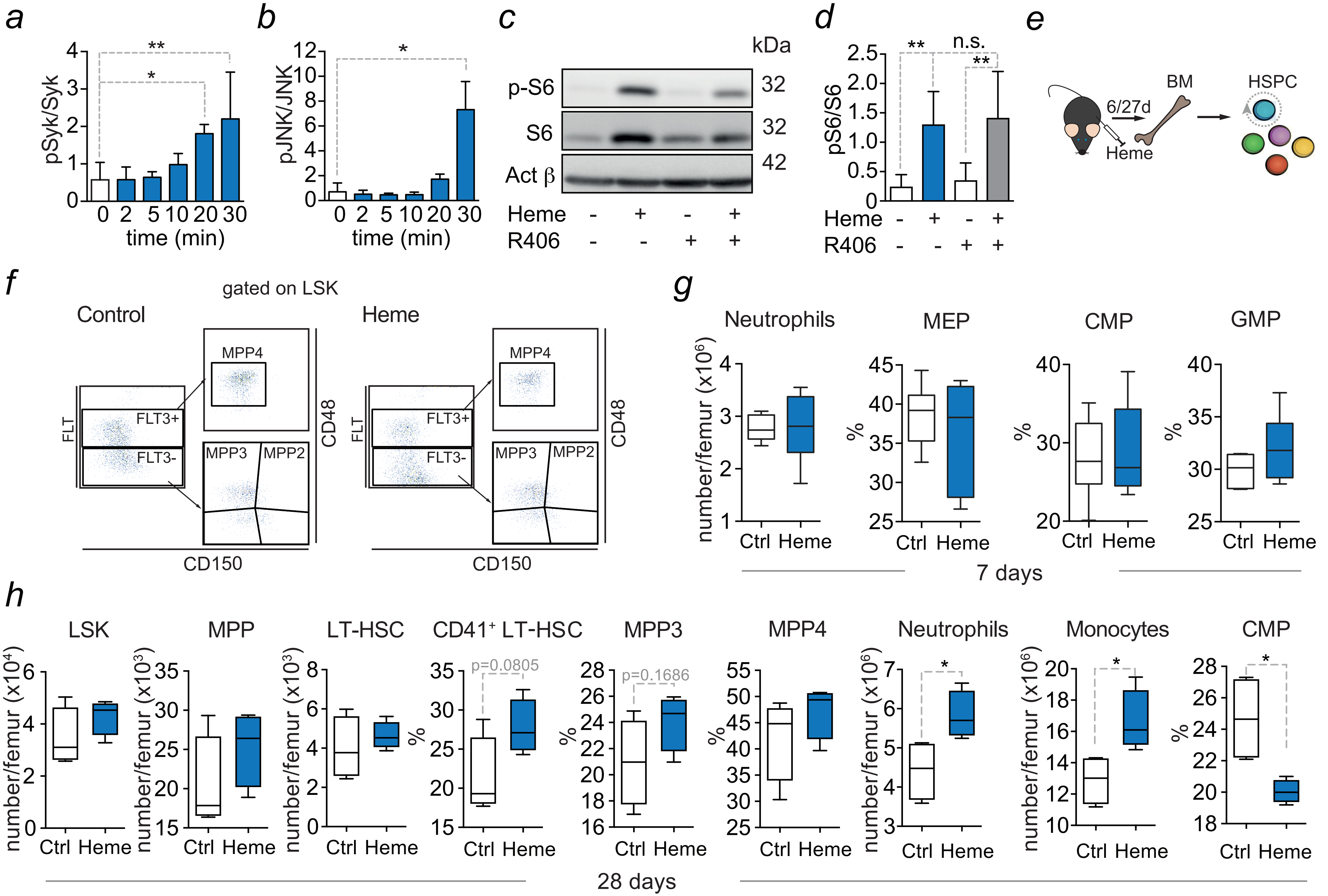

### Suppl.Fig.5

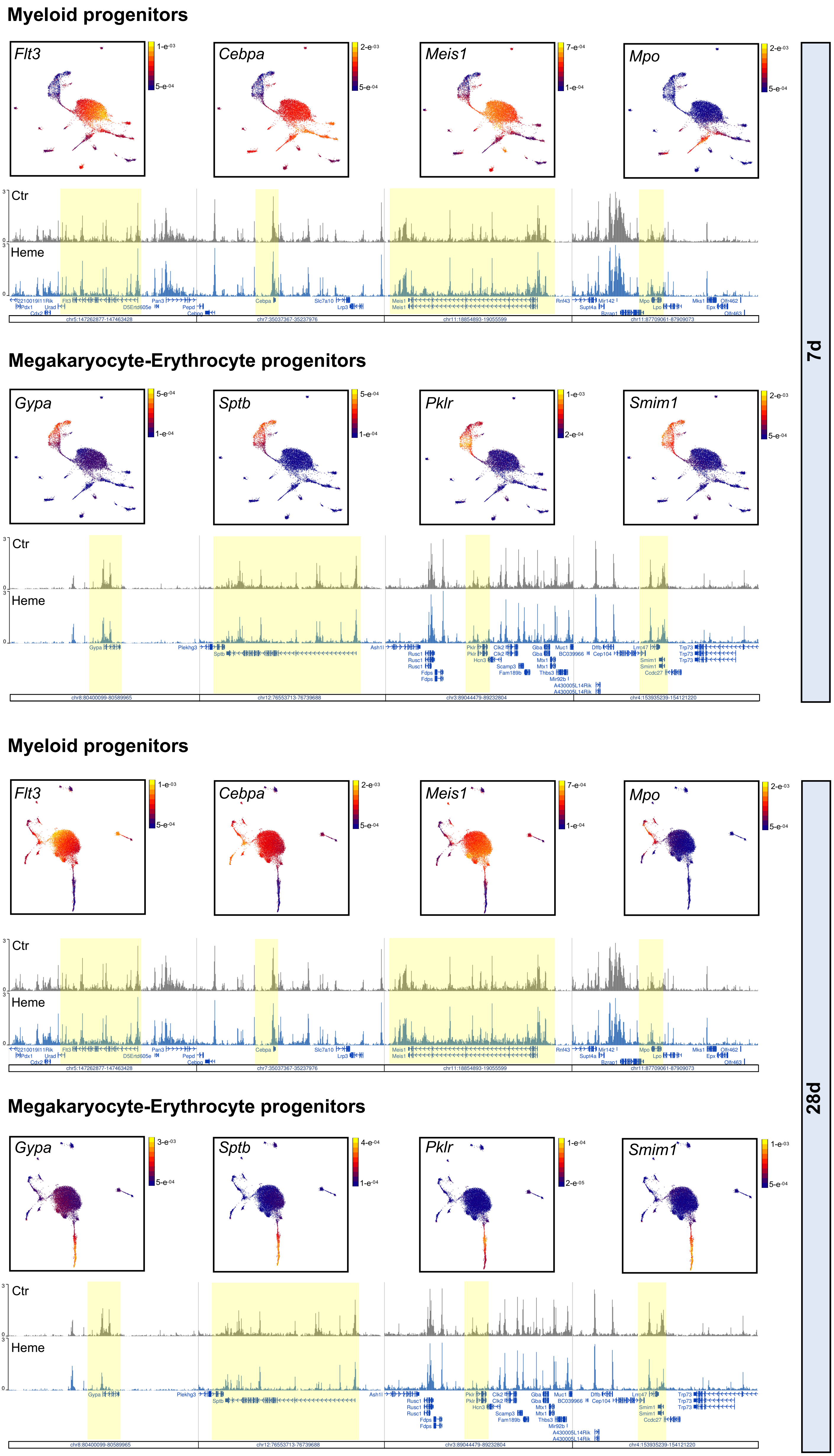

### Suppl.Fig.6

Jentho *et al.* Suppl.Fig. 4

a

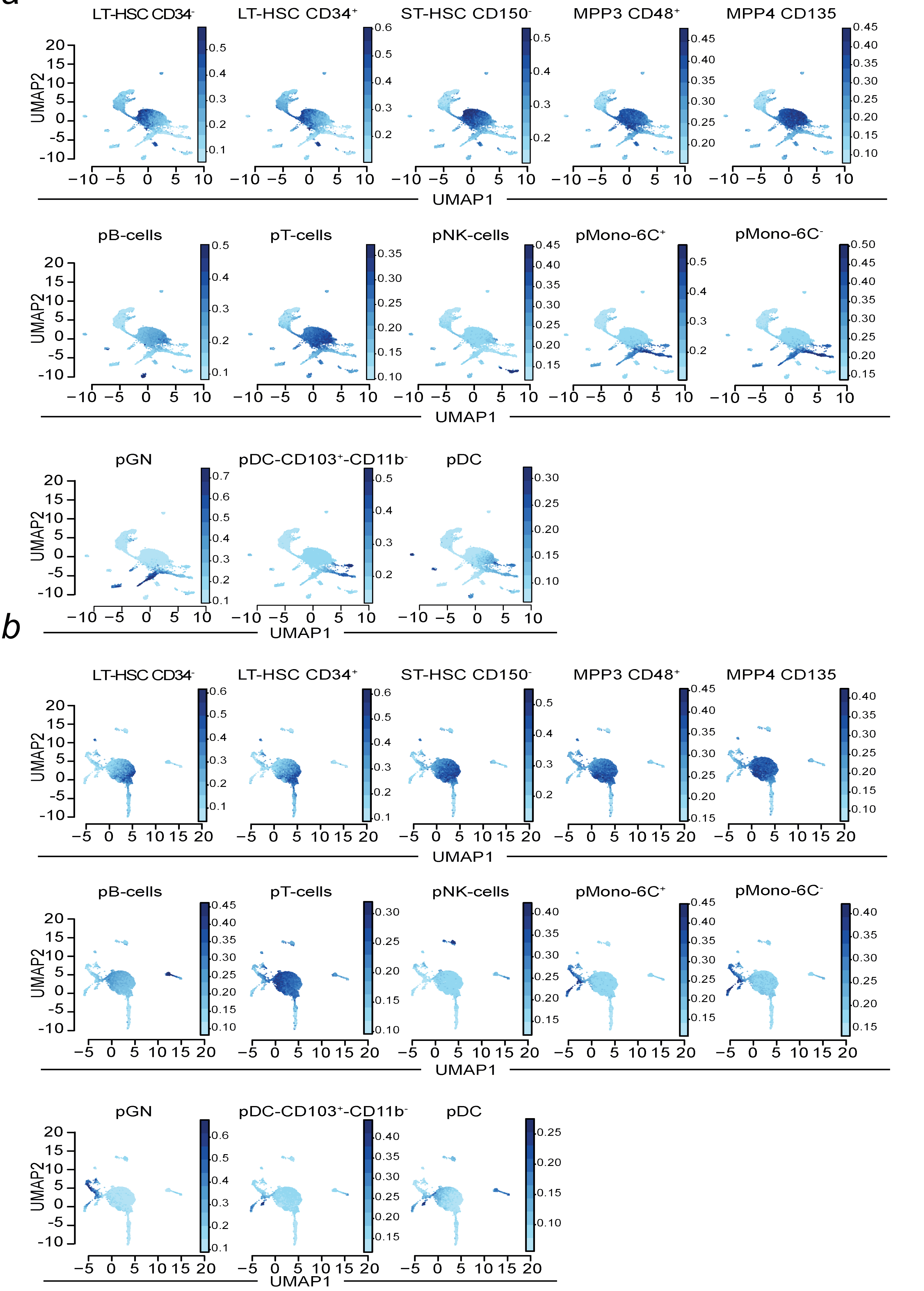
